# Differential impacts of fall versus spring prescribed burns on microbial biomass, richness, and composition in young mixed conifer forests

**DOI:** 10.64898/2026.03.09.710622

**Authors:** Basubi Binti Zhilik, M. Fabiola Pulido Barriga, Peter M Homyak, Robert A. York, Sydney I. Glassman

## Abstract

Prescribed burns aim to restore ecological processes and mitigate high-severity wildfire risks. Historically, California wildfires occur in summer or fall, but due to limited burning opportunities, prescribed burns occur in fall and spring. Yet, whether the burn season affects ecological outcomes is largely unknown. Here, we test prescribed burn season impacts on soil burn severity, bacterial and fungal abundance, richness, and composition as assessed with 16S and 18S qPCR and 16S and ITS2 Illumina MiSeq. We implemented a Before-After-Control-Impact design with 9 California montane mixed conifer forest stands (4 fall, 4 spring, 1 unburned control). We assessed resilience by collecting 6 sub-samples per stand at 6 time points ranging from pre-fire to 3 days, 1 and 6 months, and 1 and 2 years after fall and spring burns. Fall burns significantly reduced bacterial and fungal abundance and richness, whereas spring burns did not. Indeed, at 24 days after fall burns compared to pre-fire, bacterial and fungal richness were reduced by 24-30%, with richness of ectomycorrhizal fungi reduced by 45% and saprobic fungi by 28%. After the fall burn, fungal richness recovered within a year and fungal abundance within 6 months, whereas bacterial abundance and richness recovered in 2 years. Bacterial and fungal communities experienced compositional turnovers after both burns, with the emergence of several genera of “fire-loving” pyrophilous bacteria (*Massilia*, *Paenibacillus)* and fungi (*Geminibasidium, Pyronema*, *Neurospora*). Our findings suggest that, although seasonal differences were evident, both fall and spring prescribed burns mitigate wildfire impacts while promoting the succession of pyrophilous microbes.

## Introduction

Prescribed burns are controlled, low-intensity fires used to reduce surface fuels, improve stability and resilience of fire-adapted ecosystems (Bihari & Ryan, 2012; Regmi et al., 2023), and mitigate the severity of uncontrolled wildfires (Ryan et al., 2013). Managers often align burning to occur during historic wildfire seasons, which for Western North America are summer and fall (Stephens et al., 2004). Limited opportunities for burning during summer and fall due to logistical, regulatory, and weather-related constraints have prompted the use of spring prescribed burns to increase area burned (Regmi et al., 2023; Ryan et al., 2013; Striplin et al., 2020). However, it remains unclear whether burns conducted in spring produce equivalent ecological outcomes comparable to those achieved during traditional fall burn seasons, particularly for soil microbiomes, which are critical drivers of plant productivity and biogeochemical cycling (Crowther et al., 2019; Van Der Heijden et al., 2008).

Prescribed burns typically mitigate the negative impacts on soil microbiomes that are often observed after wildfires (Pressler et al., 2019). Wildfires can reduce microbial richness and biomass up to 80-90% across a variety of ecosystems (Day et al., 2020; Enright et al., 2022; Pulido-Chavez et al., 2021; Pulido-Chavez et al., 2023; Whitman et al., 2019). Alterations to microbial communities caused by wildfires can disrupt critical microbial functions, like nutrient cycling (Nave et al., 2011; Nelson et al., 2024) and plant microbe symbioses (Dove & Hart, 2017), including ectomycorrhizal fungi (EMF) and arbuscular mycorrhizal fungi (AMF), which associate with ∼80% of plant species (Brundrett & Tedersoo, 2018). Fires can also increase the abundance of pyrophilous or “fire-loving” fungi, such as *Pyronema* or *Neurospora,* or bacteria such as *Massilia* or *Bacillus*, that are absent or rare pre-fire (Enright et al., 2022; Fox et al., 2022; Whitman et al., 2019). Fire intensity, or the heat transfer of fire, and fire severity, or the impact of fire on above- or belowground organisms, can shape wildfire impacts (Keeley, 2009). Pyrophilous microbes can initiate secondary succession after both high-severity wildfires (Pulido-Chavez et al., 2023) and low-severity prescribed burns (Fischer et al., 2023). Their unique adaptations enable pyrophilous microbes to rapidly colonize and persist in fire-affected soils, where they initiate key biochemical processes crucial for the recovery of burned ecosystems (Fischer et al., 2021; Johnson et al., 2023; Semenova-Nelsen et al., 2019). However, how burning in different seasons affects microbial biomass, richness, composition, and the emergence and succession of pyrophilous microbes is largely unknown, despite the increasingly widespread practice of implementing both fall and spring prescribed burns.

While soil microbiomes are recognized as critical drivers of biogeochemistry (Crowther et al., 2019), and pyrophilous microbes drive microbial succession after fall wildfires (Pulido-Chavez et al., 2023), it remains unclear how prescribed burns in fall versus spring modulate microbial responses, including microbial resilience, or return to pre-fire conditions. Wildfires can alter soil microbiomes and lead to reduced richness for decades post-fire (Pérez-Valera et al., 2018), yet the season of the fire and timing of subsequent precipitation may affect resilience outcomes. Higher soil moisture in spring could theoretically buffer against microbial mortality (Badía et al., 2017), yet wetter soils may transfer heat more efficiently than dry soils (Vasiliev et al., 1998), leading to more microbial mortality in spring than fall burns. In Mediterranean climates, spring burns are followed by dry conditions during summer, which may further reduce microbial resilience compared to fall burns, which are typically followed by winter rains that may facilitate microbial recovery (Fajardo-Cantos et al., 2025; Hinojosa et al., 2016). Experimental burning across seasons, coupled with repeated microbial soil sampling are necessary to disentangle the effects of burn season, but these studies are rare. So far, studies have tested burn season effects on fungal communities in a Mediterranean pine forest (Vázquez-Veloso et al., 2022), on AMF in grasslands (Hopkins & Bennett, 2023), and on EMF in Eastern Mediterranean shrublands (Livne-Luzon et al., 2021) or *Pinus ponderosa* forests (Smith et al., 2004). These studies indicate that fall burns have larger impacts than spring burns on AMF traits (Hopkins & Bennett, 2023) and EMF richness (Smith et al., 2004), but that spring burns may have larger impacts on saprobic fungi (Livne-Luzon et al., 2021). Thus, while fall burns may influence mycorrhizal fungi more negatively than spring burns, more research is needed to determine how bacterial and fungal communities will respond, particularly in mixed conifer forests where both wildfires and prescribed burns tend to be more intense and severe than in grasslands or shrublands (Cox et al., 2022; Glassman et al., 2023).

Pine forests, which rely heavily on EMF for initial establishment and survival (Collier & Bidartondo, 2009; Glassman et al., 2015; Nuñez et al., 2009), cover large tracts of the terrestrial biosphere, including 65% of Western North America (Landers et al., 1995), and are a common target for prescribed burns as a management tool (Striplin et al., 2020). There is limited research indicating that spring burns lead to higher plant mortality (Greenberg’ et al., 1995; Knapp et al., 2005) and root damage (Swezy & Agee, 1991) than fall burns in mature (∼100 years old) coniferous forests. Higher mortality in spring burns is likely caused by greater plant sensitivity to heat during their period of primary growth. However, young coniferous forests (∼12-14 years old) are becoming more common due to post-fire reforestation (North et al., 2019) and may have different responses to burn season than mature forest stands (Bellows et al., 2016; York et al., 2022). Young coniferous forests in the Sierra Nevada are particularly vulnerable to high-severity fire (Levine et al., 2022) and thus have been proposed as high-priority targets for prescribed burns (York et al., 2021), but how their associated soil microbiomes respond to different burn seasons is completely unknown.

Here we asked, how do spring versus fall prescribed burns influence 1) soil burn severity and tree mortality? 2) total bacterial and fungal biomass and richness? 3) richness of fungal functional guilds, particularly EMF, that are critical coniferous tree partners? and 4) microbial composition, including the emergence and succession of pyrophilous bacteria and fungi? We hypothesized that lower soil moisture and drier conditions during fall burns would result in higher soil burn severity, leading to stronger negative effects on microbial biomass, richness, and community composition, and longer recovery time to pre-fire conditions. In contrast, higher soil moisture in spring might attenuate burn severity, reducing microbial impact. We tested our hypotheses by examining the influence of fall versus spring prescribed burns on soil burn severity, tree mortality, bacterial and fungal biomass, richness, and composition in a young mixed conifer forest (∼12-13 years old). We assessed resilience by collecting 6 sub-samples in each of 9 forest stands (4 fall, 4 spring, 1 unburned control) at 6 time points ranging from pre-fire to 3 days, 1 and 6 months, and 1 and 2 years after fall and spring burns. Our study provides insight into microbial responses to seasonal prescribed burns and provides practical guidance for selecting the most ecologically feasible burn season in fire-prone ecosystems.

## Materials and methods

### Study site

The study took place at Blodgett Forest Research Station (BFRS), which has had an experimental prescribed burn program for over 20 years (Knapp et al., 2005). BFRS is a mixed conifer forest on the western slopes of the central Sierra Nevada, California range (38 ° 54’ 45” N, 120 ° 39’ 27” W) at roughly 1320m elevation. Common tree species include Sugar pine (*Pinus lambertiana*), ponderosa pine (*Pinus ponderosa*), white fir (*Abies concolor*), incense-cedar (*Calocedrus decurrens*), Douglas-fir (*Pseudotsuga menziesii*), California black oak (*Quercus kelloggii*), tanoak (*Notholithocarpus densiflorus*) (Stephens & Moghaddas, 2005), which are all obligate EMF symbionts (Wang & Qiu, 2006). Soils at the site are classified as mesic Ultic Haploxeralfs mapped in the Holland series (Soil Survey Staff, 2024). The historical fire regime near BFRS consisted of frequent low-to-moderate severity burns, with fire return intervals from 8 to 15 years (Stephens et al., 2004). Annual precipitation (including rain and snow) is 1450 mm yr^−1^, and the mean annual temperature 6.2^◦^C with a daily maximum of 10.9^◦^C and a minimum of 2.6^◦^C (BFRS weather station data).

### Experimental Design and Prescribed Burns

Treatment Alternatives for Young Stand Resilience (TAYSR) is a long-term study evaluating several approaches for building resilience in young stands following reforestation at BFRS (Bellows et al., 2016; North et al., 2019). We selected a total of 9 forest stands within the TAYSR experiment, which we refer to as our plots. At the time of burns, plots were 12-13 years in age, 0.4 to 0.8 ha in size, and dominated by *P. lambertiana*, *P. ponderosa*, and *C. decurrens*. We burned 4 plots in the fall (11/13/2019), 4 in the spring (4/28-5/1/2020), and 1 was left unburned (control), allowing us to have a complete Before-After-Control-Impact (BACI) design (Fig. 1A), which allows for treatment effects to be distinguished from background time effects shared by all sites (Stewart-Oaten & Bence, 2001). See Table S1 for plot locations and elevations.

**Fig. 1:**
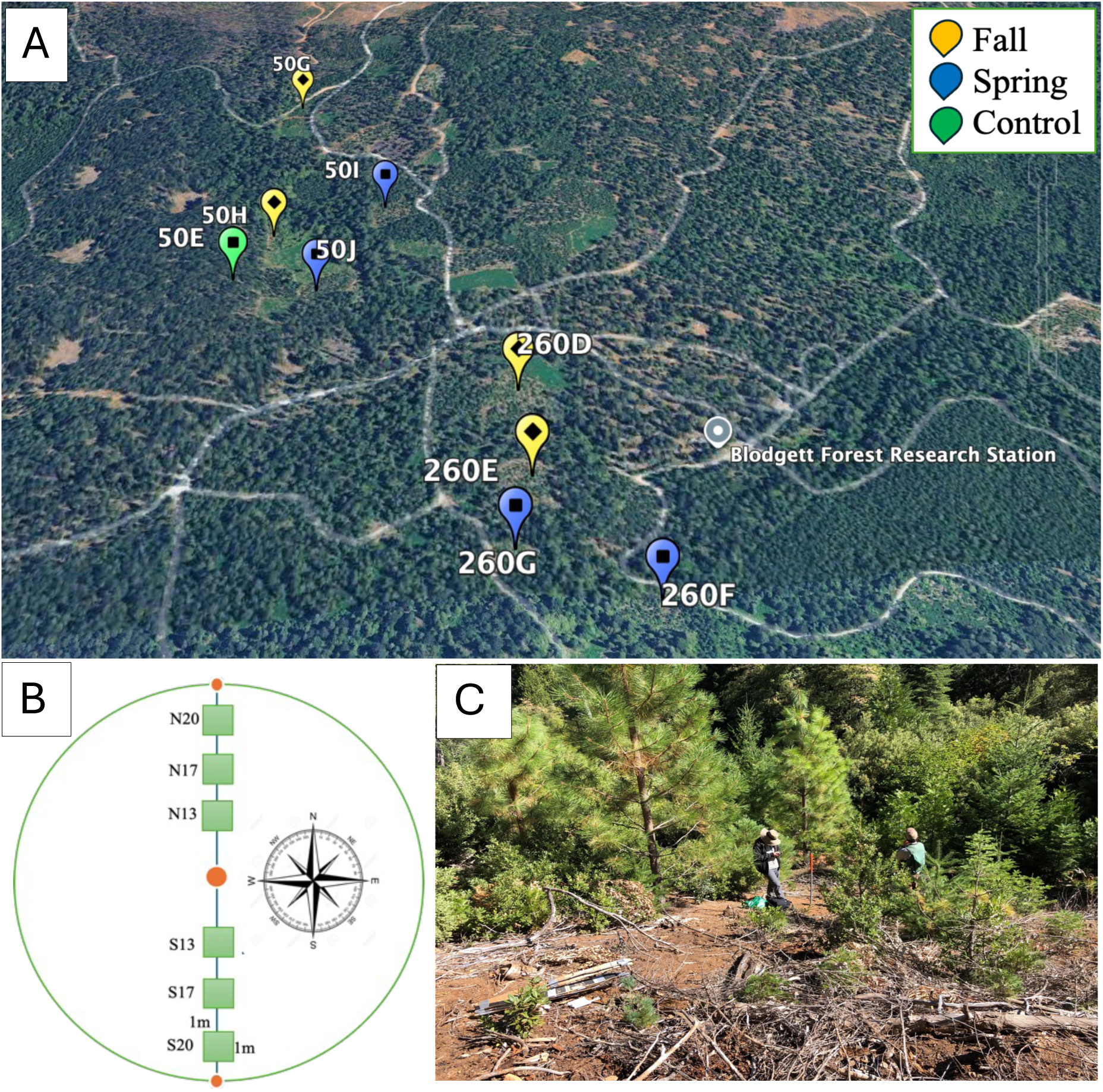
A. Stands burned in Fall 2019 (yellow) and Spring 2020 (blue), and unburned Control (green). B. Experimental design for each plot with vegetation sampling transect (blue) across the center and location of six 1m^2^ soil sampling subplots (green squares). C: Transect line with PVC pole at the BFRS.

We established plots prior to fall and spring burns on 10/03/2019, demarcating the center of each 0.4-0.8 ha stand using PVC poles (Fig. 1B). We ran two Brown’s fuel transects and a belt transect 7.3 meters wide from south to north across the stand, and we placed six 1m^2^ subplots along the transects at 13, 17, and 20 meters north and south of the plot center to allow for best congruence of plant and soil microbiome data but to not overlap with fuel measurement subplots (Fig. 1C). We tagged and measured every tree > 1.37 meter tall for species identification, height, diameter at breast height (DBH) and height to crown base (York et al., 2022).

We applied the same burn prescription for both fall and spring, defined as the range in weather conditions under which burns can occur, such that 10-hour fuel moisture was 8-9% for fall and 8-10% for spring, relative humidity was 36-40% for fall and 34-48% for spring, and air temperature was 13-17°C for fall and 12-23°C for spring. Ignitions were done with drip torches and strip-head firing, progressing slowly and representing a best management practice when the objective is to maximize surface fuel consumption while minimizing damage and mortality of the canopy (York et al., 2022). Tree mortality and crown volume scorch were measured 1-year post-burn (York et al., 2022). Using a ruler, we measured and averaged ash depth from 3 locations per 1m^2^ subplot at 3 days post-fire as a proxy for soil burn severity as previously established (Glassman et al., 2023; Pulido-Chavez et al., 2023).

### Soil Collection

We collected soils from 6 subplots from all 9 plots pre-fire on 10/03/2019, and post-fire at 3 days, 24 days, 6 months, and 1 and 2 years for fall burns and 3 and 24 days, 1, 1.5, and 2 years for spring burns since collection at 6 months after the spring burn was not feasible due to deep snow that prevented entry to the sites. Soils from the unburned control plot were collected at each post-fire time point for both seasons. We used a releasable bulb planter (cleaned with 70% ethanol between samples) to collect two 10-cm deep soil cores per 1 m^2^ subplot. Soil samples were stored on ice and shipped to the Glassman lab at the University of California, Riverside (UCR), within 48 hours, where we homogenized soil samples, stored 20-30 g at -80 °C for molecular work, measured soil gravimetric moisture content from 5-10 g, and air-dried the rest for analysis of carbon cycling (Krichels et al., 2026). Overall, we collected 354 soil samples (288 = 8 plots x 6 samples/plot x 6 time points) + 54 unburned control samples (6 samples/1 plot x 9 time points). Note that the control plot was sampled 9 times rather than 12 because 3 timepoints overlapped. For example, the pre-fire timepoint was shared by all plots, and 6 months after the fall burn fell on the same day as 3 days after the spring burn.

### DNA extractions, PCRs, and sequencing

We used Qiagen DNeasy Power SoilPro Kits (Qiagen, Maryland, USA) to extract DNA from 0.25g of each frozen soil sample following the manufacturer’s protocol with the modification at C1 such that 100 µl of ATL solution was added to 700µl of C1 solution and incubated overnight at 4°C to improve DNA yields. DNA extractions were PCR-amplified via two-step PCR reactions, the first to amplify the regions of interest and the second to ligate the barcodes and adapters following the Dual-Index Sequencing Strategy (DIP) (Kozich et al., 2013). We amplified the prokaryotic V4 region of the 16S rRNA gene with primer pair 515F and 806R (Caporaso et al., 2011) and fungal Internal Transcribed Spacer 2 (ITS2) with the primer pair ITS4-fun and 5.8s (Taylor et al., 2016). Although both archaea and bacteria were detected by the 16S primers, we simplify terminology by referring to the results solely as bacterial, since archaea accounted for only 0.4% of the sequencing reads.

For PCR1, we combined 5 μl of undiluted DNA for fungi and 1 μl of 1:10 diluted DNA for bacteria, 0.5 μl of each gene-specific primer (10 μM), 12.5 μl of AccuStart II PCR Tough Mix (Quantabio, Beverly, MA, USA), and Ultra-Pure Sterile Molecular Biology Grade water (Genesee Scientific, San Diego, CA, USA) to complete the 25 µl reaction. Thermocycler settings were as follows: 94°C for 2 min, followed by 29 (bacteria) or 30 (fungi) cycles of 94°C for 30s, 55°C for 30s, 68°C for 2 min, ending with a 10 min extension at 68°C before a 4°C hold. We always included negative DNA extraction and PCR controls. We visualized PCR products with gel electrophoresis to ensure amplification and cleaned products with the AMPure XP magnetic Bead protocol (Beckman Coulter Inc., Brea, CA, USA). For PCR2, DIP primers containing the barcodes and adaptors were ligated to the amplicons in a 25 μl reaction containing 2.5 μl of the 10 mM DIP PCR2 primers, 6.5 μl of ultrapure water, 12.5 μl of Accustart II PCR ToughMix, and 1 μl of 1:10 diluted PCR1 product. Thermocycler conditions for PCR2 were 94^◦^C for 2 min followed by 8 cycles of 94°C for 30s, 60°C for 30s, 72°C for 1 min, and ending at a 4°C hold. Bacterial and fungal PCR products were then separately pooled based on gel electrophoresis band strength and cleaned with AMPure following established methods (Glassman et al., 2023; Pulido-Chavez et al., 2023). The 16S and ITS pools were each checked for quality and quantity with an Agilent 2100 Bioanalyzer, then pooled at 0.4 bacteria to 0.6 fungi ratio prior to sequencing with Illumina MiSeq 2x300 bp at the UCR Institute for Integrative Genome Biology. To contain all samples, we ran 2 Illumina MiSeq runs, each containing up to 192 samples for bacteria and fungi. Sequences are publicly available at the NCBI SRA under accession number PRJNA1417007.

### Bacterial and fungal abundance

We estimated gene copy number of 18S rRNA using FungiQuant primers (Liu et al., 2012) and of 16S rRNA using Eub338/Eub518 primers (Fierer et al., 2005) to estimate fungal and bacterial abundance as a proxy of biomass. The qPCR master mix was prepared with 5 μL of 1X iTaq SYBR mix (Bio-Rad, Hercules, CA) and 0.4 μl of 10µM of each primer, with 1 μl of DNA and ultrapure water for a final volume of 10 μL. Each reaction was run in triplicate on a BioRad CFX Opus 384 Real-Time PCR System starting at 94^◦^C for 5 min, followed by 40 cycles of a denaturing step at 94^◦^C for 20s, primer annealing at 52^◦^C for bacteria or at 50^◦^C for fungi for 30s, and an extension step at 72^◦^C for 30s. Standards were generated (Pulido-Chavez et al., 2023), and gene copy numbers per gram of soil were calculated (Joukhajian et al., 2026) as previously established.

### Bioinformatics

We analyzed paired-end demultiplexed Illumina sequence files using QIIME2 version 2020.8 pipelines (Bolyen et al., 2019). The sequencing facility removed the 5′ primers and adapter sequences, and we removed the 3′ primers and adapters using the Qiime2 CutAdapt plugin (Martin, 2011). We used DADA2 to filter out and remove chimeric sequences and low-quality regions and produce Amplicon Sequence Variants (ASVs) (Callahan et al., 2016). For taxonomic analysis, we used the fitted classifier with the classify-sklearn plugin (Pedregosa et al., 2011) and aligned fungal reads against the UNITE classifier v8.2 (Abarenkov et al., 2010) and bacterial reads against the Silva 16S classifier 2020 v132 (Quast et al., 2012). We removed the mitochondrial and chloroplast reads for bacteria, and any fungal reads that did not match to Kingdom Fungi. We also specified fungal guilds with FUNGuild (Nguyen et al., 2016) and retained only guilds classified with a highly probable confidence level assignment.

### Statistical analysis

All statistical analyses were conducted in R v4.2.0 (R Core Team, 2022) with figures made in ggplot2 (Wickham, 2016), and all scripts to generate statistics and figures can be found at: https://github.com/Basubi1/Young-Blodgett-Rscipt. Our Illumina MiSeq runs resulted in 13.9M bacterial and 22.4M fungal reads, and 34,348 bacterial and 16,500 fungal ASVs. We estimated microbial richness as the observed number of ASVs after rarefying all samples to the same number of sequences (4,012 for bacteria, 5,132 for fungi) following best current practices for microbiome analysis (Schloss, 2024). We tested data for normality with the Shapiro-Wilk test and heteroscedasticity by graphical inference via the residuals. For all response variables where the conditions of normality could not be met, we employed generalized linear mixed-effects models (GLMER) using the lme4 package (Johnson et al., 2015). To answer Q1, we used analysis of variance (ANOVA) on tree mortality and ash depth, followed by a post-hoc Tukey test to determine which burn season had a larger impact on soil burn severity. To answer Q2 and Q3, we used GLMER to test the impacts of fire, time, and their interactions in control, fall, and spring plots individually against the pre-fire timepoint. Response variables included total bacterial and fungal species biomass and richness, and richness of ectomycorrhizal, saprobic, and plant pathogenic fungi. All models were run independently. We used backward elimination using the Akaike Information Criterion (AIC) (Burnham & Anderson, 2004) and tested the significance of the best-fit model with ANOVA (p < 0.01) to select the best model using the MASS package (Carmona, 2004). Considering overdispersion and conditional variance higher than the mean, we selected a negative binomial distribution (Bliss & Fisher, 1953; Ross & Preece, 1985). We included time, plot, and subplot as random effects, based on comparisons with a null model using AIC. We calculated marginal and conditional R^2^ values using the package MuMIn Barton.

To answer Q4, we tested the impacts of burn season and time on bacterial or fungal community composition by creating square root transformed Bray-Curtis Dissimilarity matrices with Vegan’s avgdist function and using betadisper to test the homogeneity of group dispersion (Oksanen et al., 2001). We used PERMANOVA (Anderson, 2017) as implemented in vegan’s adonis2 function (Oksanen et al., 2001) with 999 permutations and including time as a random effect. Employing previously used techniques (Pulido-Chavez et al., 2023), we tested the succession of bacterial and fungal community composition and counted how many times the community turned over by calculating Euclidean distance between time points using the Vegan package and using a Principal Coordinate Analysis (PCoA) to visualize the centroid and standard error between time points. We considered each time point that did not have overlapping standard error bars in either the x or y directions to be a turnover. We fit a negative binomial distribution using the DESeq2 package (Love et al., 2014) to identify the pyrophilous taxa with significant positive or negative log2fold changes at each post-fire time point compared to pre-fire conditions. DESeq results were summarized into a figure by plotting the log2-fold change values for each taxon that significantly responded to fire at each time point for both seasons, as previously published (Joukhajian et al., 2026). We further visualized changes in the community and pyrophilous microbes by creating relative sequence abundance bar plots of the dominant bacterial and fungal phyla and genera using phyloseq v1.24.2 (McMurdie & Holmes, 2013).

## Results

### 3.1 Effect of burn season on soil burn severity, tree mortality, and soil moisture

Both fall and spring burns resulted in tree damage that would translate to a low to moderate severity fire (tree mortality of 35.8% ± 7 for fall and 28.5% ± 4 for spring; Fig. S1A), with fall tending to have higher soil burn severity than spring burns, where average ash depth at 3 days post-fire was 1.95 ± 0.4 cm for fall compared to 0.83 ± 0.1 cm for spring (Fig. S1B). Although tree mortality was not significantly higher in fall compared to spring burns, fall burns lowered surface fuel loads by 80% compared to 30% in spring (York et al., 2022). Soil moisture, which ranged from 31 to 55%, was the same in all plots at pre-fire, but was higher in the unburned control than burned plots after both fall and spring burns, which did not differ by burn season (Fig. S2, Table S2). Precipitation, which was significantly and positively correlated with soil moisture (Pearson r = 0.46, P < 0.05), peaked in December 2019 and in April 2022, corresponding to 24 days after fall and 2 years after spring burn (Fig. S3), leading to significantly increased soil moisture for both burned and control plots at 24 days after the fall burn, and in the control plot at 2 years after the spring burn (Fig. S2).

### 3.2 Effect of prescribed burn season on soil bacterial and fungal richness

Both bacterial (Table S3) and fungal richness (Table S4) were significantly reduced by fall burns and were unaffected by spring burns (Fig. 2). After fall burns, soil burn severity was significantly and negatively correlated with bacterial (R=-0.48, p <0.0001; Fig. S4A) and fungal richness (R=-0.63, p <0.0001; Fig. S4C). However, after the spring burns, there was a weaker correlation with soil burn severity for richness of fungi (R=-0.32, p <0.001; Fig. S4D) and none for bacteria (Fig. S4B). At 24 days after fall burns, richness was reduced from pre-fire conditions by 30% for bacteria (Table S5) and 24% for fungi (Table S6). Fungi were more resilient than bacteria, with richness recovering after fall burns by 1 year for fungi (Fig. 2C) and by 2 years for bacteria (Fig. 2A). Meanwhile, richness increased temporarily at 3 days for bacteria (Fig. 2A) and at 2 years for fungi (Fig. 2C) in the control plot after fall burns. In contrast, richness of bacteria (Fig. 2B) and fungi (Fid. 2D) remained unchanged in all plots after spring burns. There was no clear correlation between precipitation and richness of bacteria (Table S3) or fungi (Table S4), suggesting that richness effects were caused by burns and not precipitation.

**Fig. 2:**
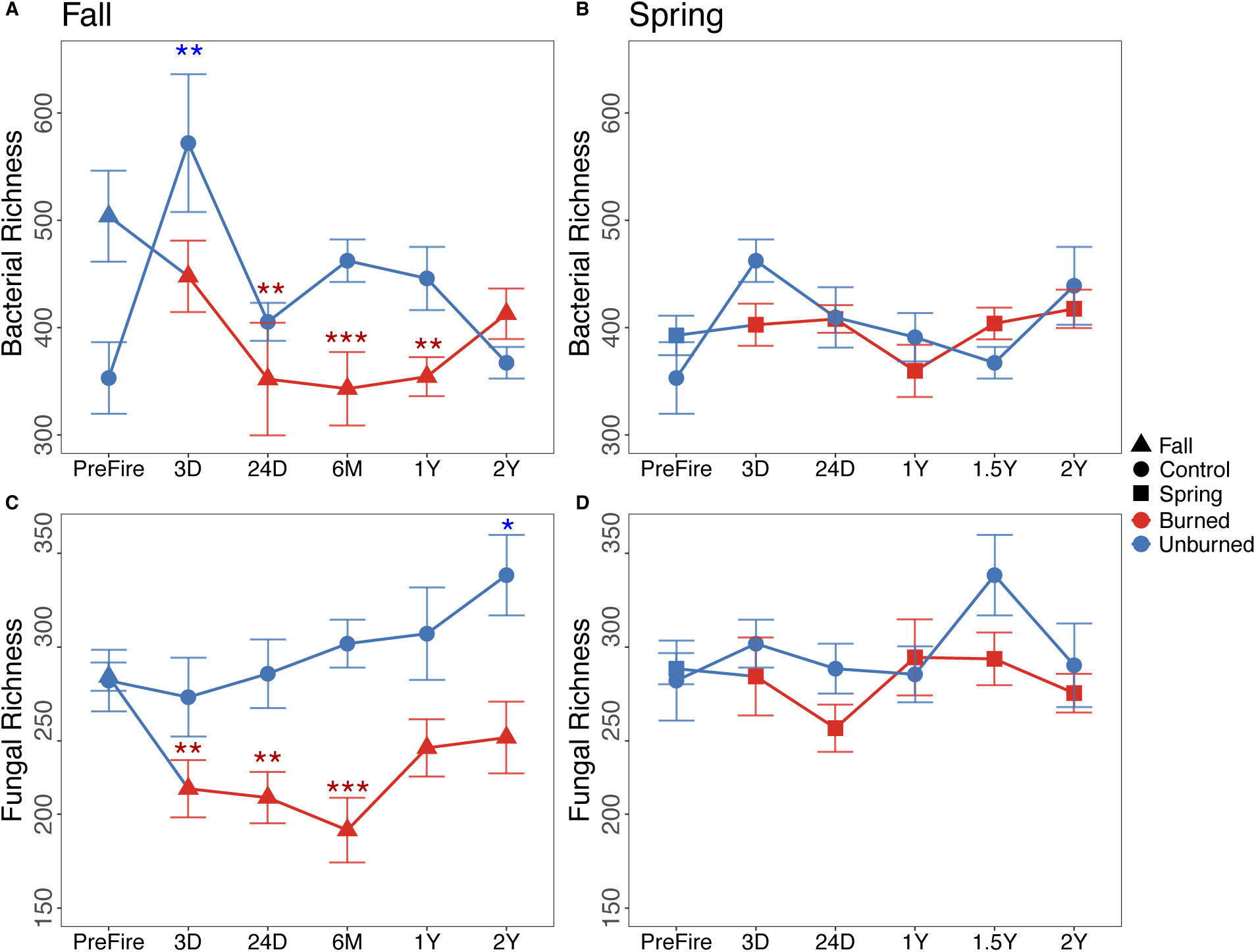
Mean richness per plot plus or minus standard error for A) bacteria in fall, B) bacteria in spring, C) fungi in fall, and D) fungi in spring. Colors indicate unburned plots from pre-fire and control (blue) versus post-fire burned (red). Shapes indicate different fall (triangle; 4 plots), spring (square; 4 plots), and control (circle; 1 plot) plots, with each plot having 6 subplots per plot. Asterisks (‘***’ P < 0.001, ‘**’ P < 0.01) indicate significant differences at each time point compared to pre-fire for either unburned (blue asterisks) or burned (red asterisks) plots. X axis represents either pre-fire or days (D), months (M), or years (Y) post-fire.

Like total fungal richness, EMF and saprobic fungal richness decreased after fall and were unaffected by spring burns (Fig. 3). Richness of EMF was reduced by 45% and saprobic fungi by 28% at 24 days after fall burn compared to pre-fire (Table S7), but richness of saprobic fungi recovered by 1 year and EMF by 2 years (Fig. 3). In contrast, fungal pathogen richness was reduced temporarily by both fall and spring burns, with a significant dip in richness at 6 months after fall and 24 days after the spring burn relative to pre-fire (Table S7). In control plots, EMF richness did not change over time, but the richness of fungal saprobes and pathogens increased at 1.5 years after spring burns, and fungal pathogen richness increased at 2 years after fall burns.

**Fig. 3:**
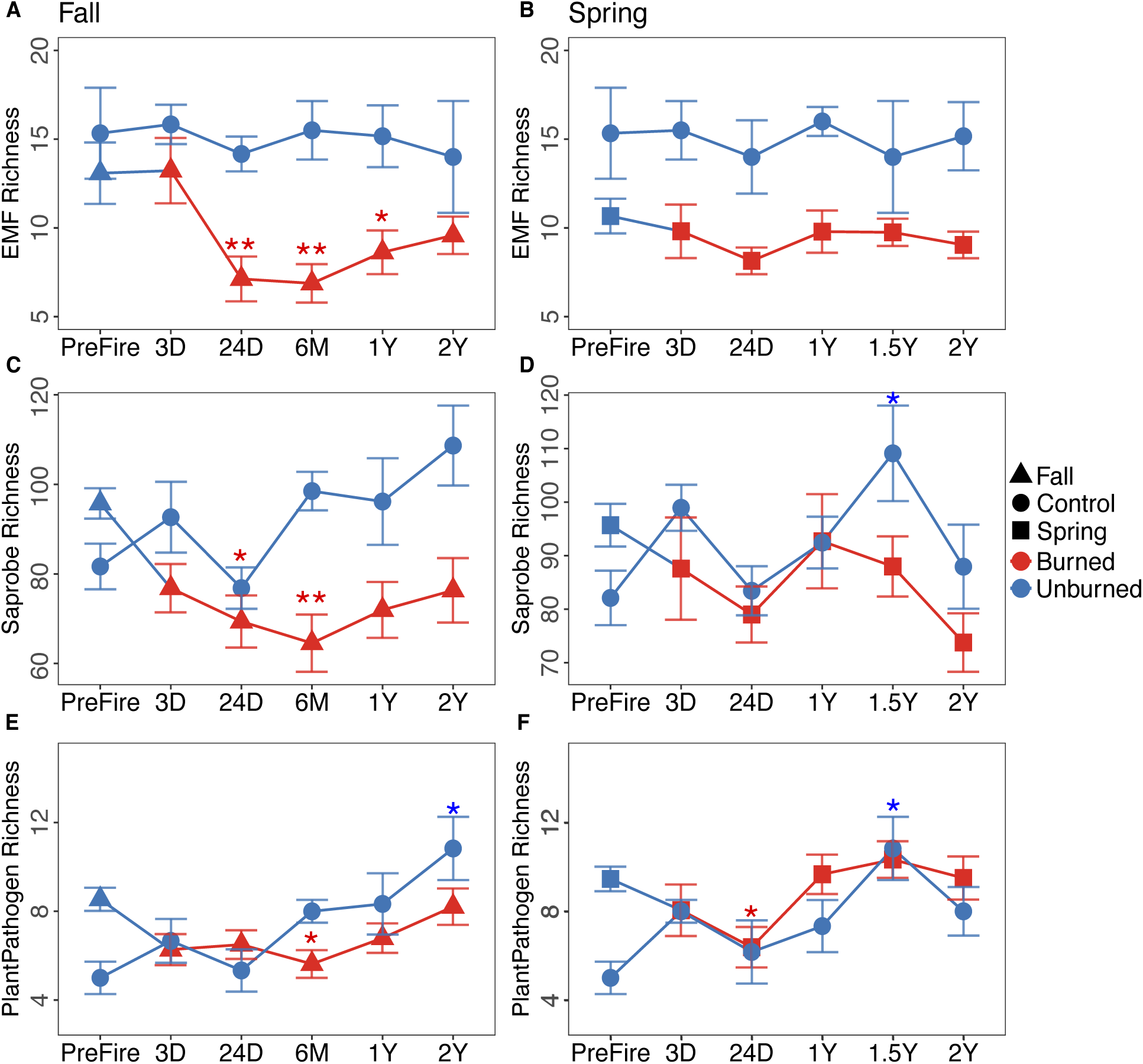
Per plot mean plus or minus standard error richness for fungal guilds of A) EMF in fall, B) EMF in spring, C) saprobes in fall, and D) saprobes in spring, E) plant pathogens in fall an F) plant pathogens in spring. Colors indicate unburned plots from pre-fire and control (blue) versus post-fire burned (red). Shapes indicate different fall (triangle), spring (square), and control (circle) plots. Asterisks (‘**’ P< 0.01, ‘*’ P< 0.05) indicate significant differences at each time point compared to pre-fire for either unburned (blue asterisks) or burned (red asterisks) plots. X axis represents either pre-fire or days (D), months (M), or years (Y) post-fire.

### 3.3 Effect of burn season on soil microbial abundance

Like richness, both 16S and 18S gene copy number (proxy of bacterial and fungal biomass) reduced following the fall burn but remained unaffected by the spring burn (Fig. 4). Compared to pre-fire, after the fall burn, bacterial biomass declined by 60% at 3 days, and 72% at 6 months, but recovered by 2 years (Fig. 4A), and fungal biomass declined by 50% at 3 days, and 65% at 24 days, but recovered by 6 months (Fig. 4B). Soil burn severity was significantly negatively correlated with biomass of bacteria (R=-0.23, p <0.01; Fig. S5A) but not with fungi after fall burns (Fig. S5C), or with bacteria (Fig. 5B) or fungi after spring burns (Fig. S5D). In contrast, in the unburned control plot, both bacterial and fungal biomass remained unchanged compared to pre-fire levels except for a temporary increase in fungal biomass 24 days after the fall burn (Fig 4C), corresponding to the peak soil moisture (Fig. S2) and precipitation (Fig. S3A).

**Fig. 4:**
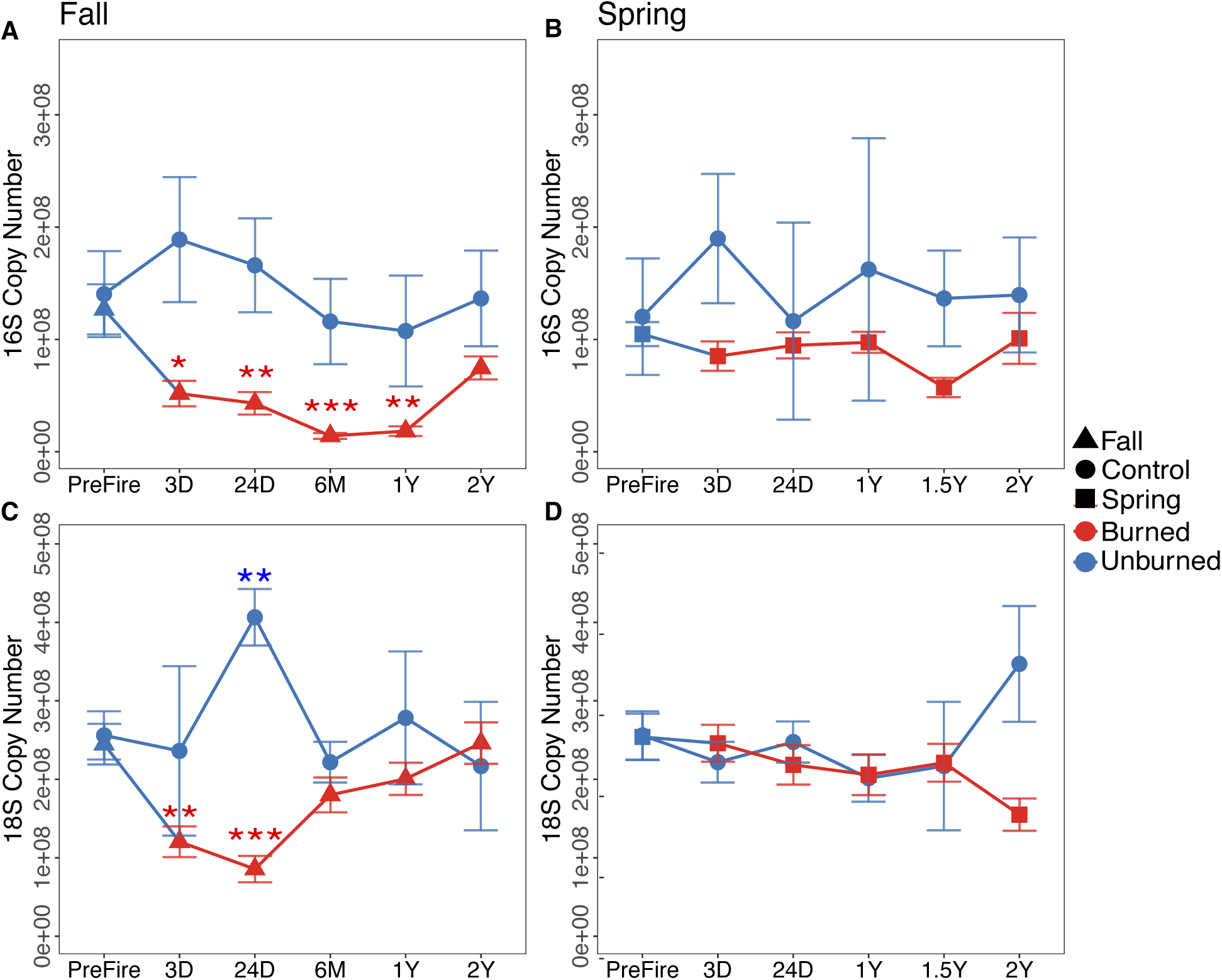
Per plot mean plus or minus standard error estimated biomass in terms of estimated gene copy number of A) 16S in fall, B) 16S in spring, C) 18S in fall, and D) 18S in spring. Colors indicate unburned plots from pre-fire and control (blue) versus post-fire burned (red). Shapes indicate different fall (triangle), spring (square), and control (circle) plots. Asterisks (‘***’ P < 0.001, ‘**’ P< 0.01, ‘*’ P< 0.05) indicate significant differences at each time point compared to pre-fire for either unburned (blue asterisks) or burned (red asterisks) plots. X axis represents either pre-fire or days (D), months (M), or years (Y) post-fire.

**Fig. 5:**
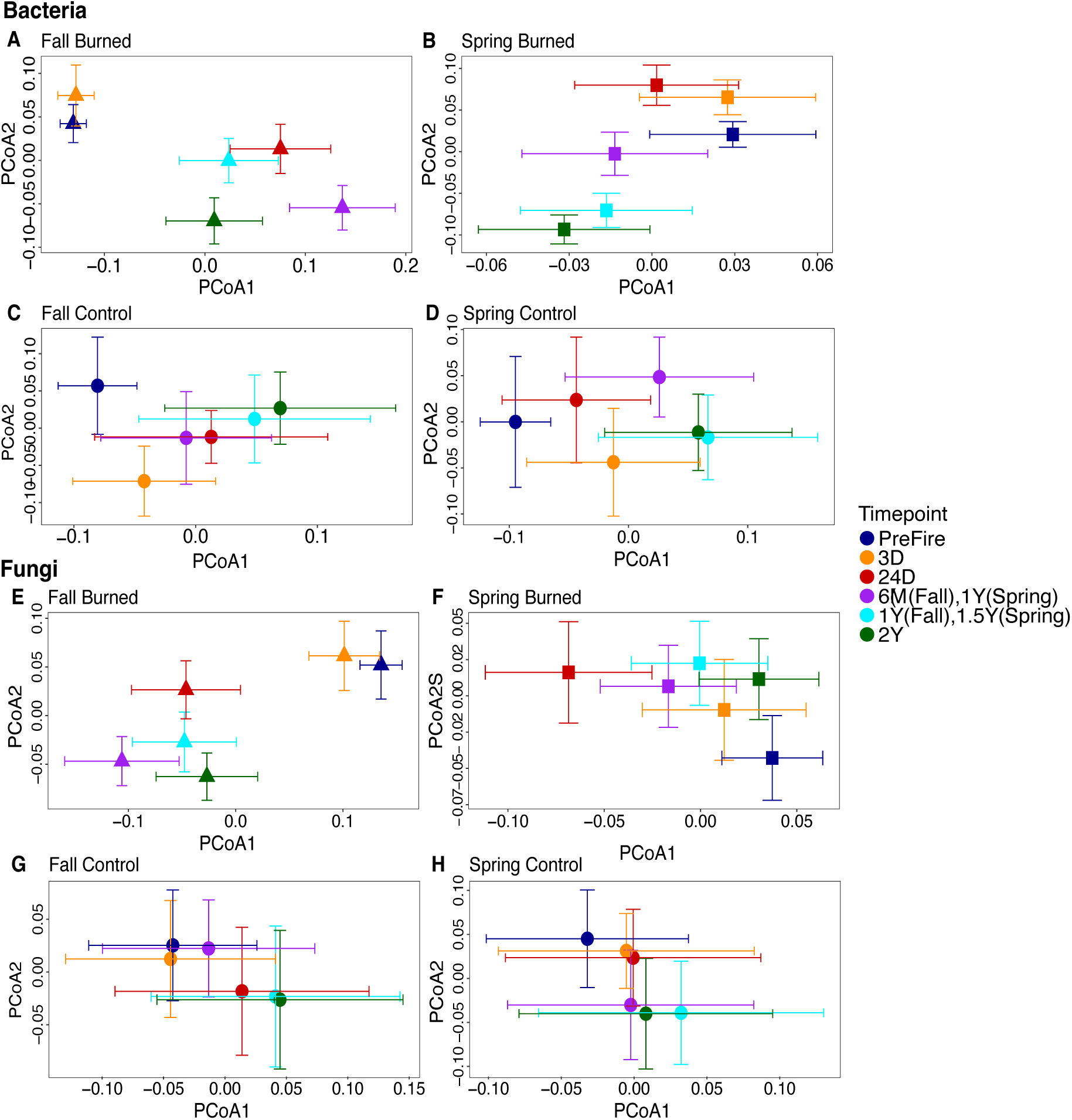
Principal Coordinates Analysis (PCoA) of mean and standard error Bacterial Bray-Curtis dissimilarity at each time point. A significant turnover in community composition is considered if error bars do not overlap. Community composition for bacteria for burned plots in A) fall and spring) and control plots in C) fall and D) spring. Community composition for fungi in burned plots in E) fall and F) spring, and control plots in G) fall and H) spring. Shapes indicate different fall (triangle), spring (square), and control (circle) plots. Colors indicate different timepoints as either pre-fire or days (D), months (M), or years (Y) post-fire.

### 3.4 Effect of prescribed burn season on bacterial and fungal community turnover

There were significant fire, time and fire by time interaction effects for both bacteria and fungi (Table S8) such that bacterial and fungal communities changed in response to fire and over time after fall and spring burns (Fig. 5). Neither bacteria nor fungi returned to pre-fire composition by 2 years after fall (Fig. 5A, 5E) or spring burns (Fig. 5B, 5F).

For bacteria, fire explained 2-3% of the variance in community composition after fall and spring burns, whereas time and fire by time interactions explained 8-9% of variance after both burns compared to pre-fire (Table S8). After fall burns, bacterial communities turned over at 6 months, 1 year and 2 years in the control plot (Fig. 5C), where in burned plots, they turned over at each time point (Fig. 5A, Table S9). In spring, burned bacterial communities also turned over at each time point (Fig. 5B, Table S9), whereas unburned communities were stable over time, except for 3 days and 1 year post-fire (Fig. 5D).

For fungi, fire explained 2-3% of the variance in community composition after fall and spring burns, whereas time explained 10-11%, and fire by time interactions explained 15% of the variance after both burns (Table S8). Fungal communities showed large turnovers at 24 days, 6 months, 1 year, and 2 years after fall burns (Fig. 5E) and at each time point after spring burns (Fig. 5F, Table S9). In contrast, fungal communities in the unburned control plot turned over at 3 and 24 days, and 2 years after fall (Fig. 5G) and at 2 years after spring burns (Fig. 5H).

### 3.5 Effect of prescribed burn on the emergence and succession of pyrophilous microbes

Bacterial communities were dominated by the phyla Actinobacteria and Proteobacteria, followed by Acidobacteria in all the burned and control plots for both seasons (Fig. 6A-D). In the burned plots, Verrucomicrobiota disappeared, and Firmicutes were abundant at all time points for both fall (Fig. 6A) and spring burns (Fig. 6B). Firmicutes were particularly affected by the fall burn, where they temporarily increased from pre-fire levels by 150% at 3 days post-fire (Fig. 6A). The Firmicutes genus *Paenibacillus* temporarily dominated 3 days after the spring burn (Fig. 6F) and increased in abundance from 3 days to 1 year after fall burns (Fig. 6E). Notably, the Proteobacteria genus *Massilia* increased from pre-fire levels by 200% at 24 days after the fall burn. *Massilia* was abundant through 2 years after the fall (Fig. 6E) and 1.5 years after the spring burn (Fig. 6F). In contrast, *Massilia* was not abundant in control plots at any time point after spring (Fig. 6H) or at 6 months through 2 years after fall burns (Fig. 6G). Two genera within the Actinobacteria order Vicinamibacterales, 67-14 and genus *Conexibacter*, were dominant at most time points in control plots (Fig. 6G, H) and after the spring burn (Fig. 6F), but disappeared by 24 days after the fall burn (Fig. 6E).

**Fig. 6.**
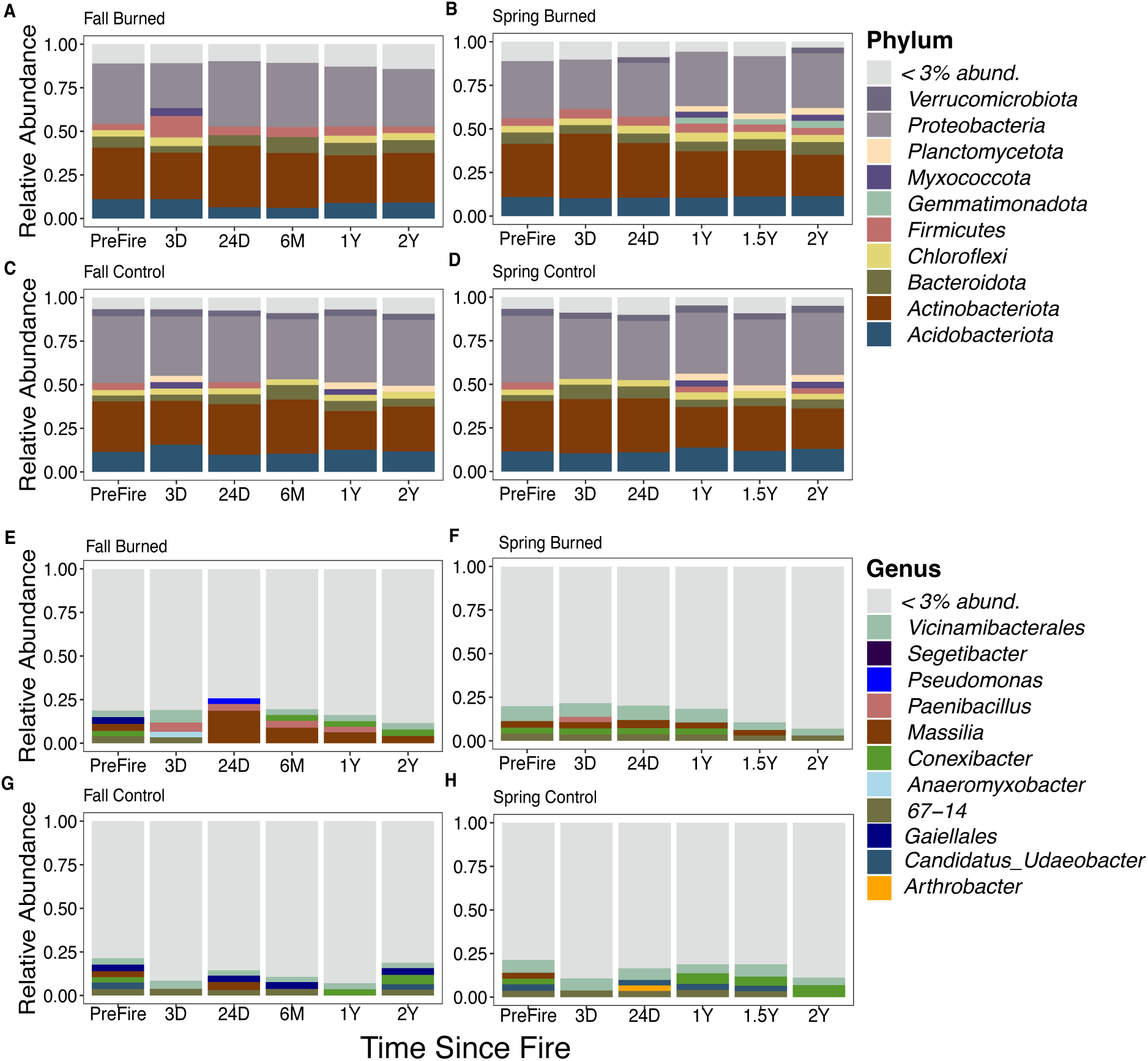
Taxonomy bar pots showing relative percent sequence abundance of bacteria- Phylum (A. fall burned B. spring burned C. fall control and D. spring control plots) and Genus (E. fall burned F. spring burned G. fall control and H. spring control plots) at all time points from pre-fire to 3 and 24 days (D) and 6 months (M) and 1,1.5 or 2 years (Y) after fire. All genera less than 3% abundant are condensed to improve visualization.

Fungal communities were largely dominated by the phyla Ascomycota and Basidiomycota in all plots for both seasons (Fig. 7A-D), however, Basidiomycota increased compared to pre-fire after prescribed burns in fall (Fig. 7A) but not spring (Fig. 7B). The increase in Basidiomycota relative to pre-fire after the fall burn was largely driven by increases in Basidiomycete yeasts, including the genera *Geminibasidium*, which increased by 266% at 3 days, and *Curvibasidium* and *Rhodosporidiobolus,* which were temporarily abundant at 24 days (Fig. 7E), and *Naganishia* and *Leucospiridium,* which became abundant at 24 days and persisted through 2 years after the fall burn (Fig. S6). In addition, the EMF Basidiomycota genus *Rhizopogon* temporarily increased by 431% at 3 days post-burn. For Ascomycota, some taxa that were cryptic or rare in the unburned plot (Fig. 7G), such as *Calypotrozyma* and *Venturia,* were abundant after the fall burn from 6 months to 2 years (Fig. 7E). In contrast to the fall burn, most of the dominant taxa remained stable after the spring burn (Fig. 7F) except for the notable increase in relative abundance of the pyrophilous Ascomycete genera *Neurospora* at 3 and 24 days and *Pyronema* at 24 days (Fig. S6). Some taxa, such as *Chalara* and *Hormonema,* were present only in the control plot (Fig. 7H), whereas *Penicillium*, *Umbelopsis*, *Odiodendron, Cladophialophora,* and *Pseudeurotium* were consistently abundant in both burned and control plots for both seasons (Fig. 7E, F, G, H).

**Fig. 7:**
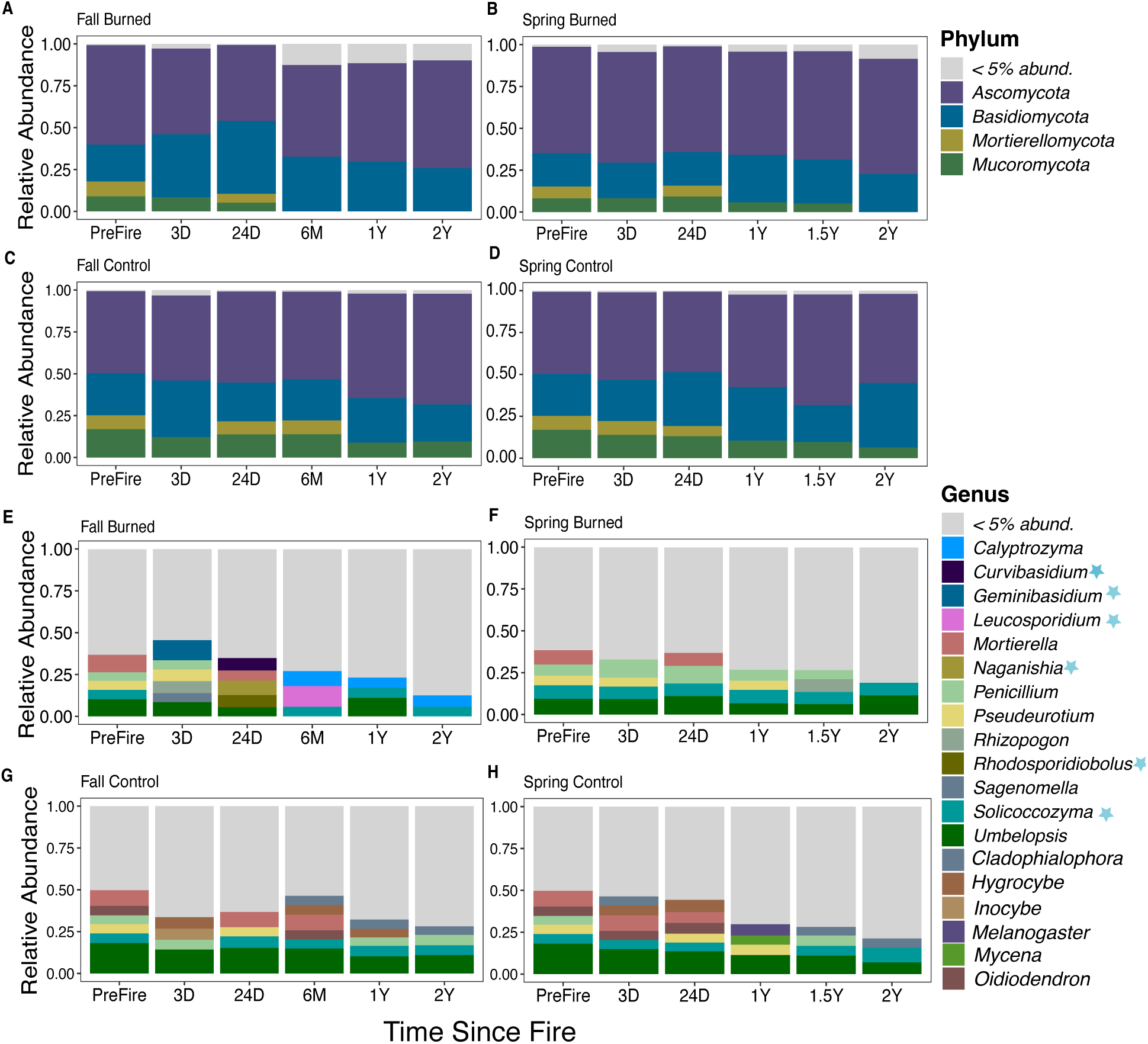
Taxonomy barplots showing relative percent sequence abundance of fungi- Phylum (A. fall burned B. spring burned C. fall control and D. spring control plots) and Genus (E. fall burned F. spring burned G. fall control and H. spring control plots) at all time points from pre-fire to 3 and 24 days (D) and 6 months (M) and 1,1.5 or 2 years (Y) after fire. All genera less than 5% abundant are condensed to improve visualization. Blue asterisks on legends indicate yeasts.

For bacteria, when testing significant changes in log2fold abundances in burned plots compared to pre-fire, 4 taxa (*Massilia*, *Pseudomonas, Tardiphaga,* and *Segetibacter*) in the fall and 3 taxa (*Conexibacter, Pedosphaeraceae*, and *Anaeromyxobacter*) in the spring showed positive changes (Fig. 8A) whereas 5 taxa (*Puia, Blastococcus, WD260*, *Arthobacter,* and *Massilia*) in fall burn and 4 taxa (*Blastococcus*, *Arthobacter, Sphigomonas,* and *Massilia*) in spring burn responded negatively (Fig. 8B). Notably, *Massilia* showed 4.5X log2fold changes compared to pre-fire at 24 days after fall but not spring burn. *Massilia* then increased to a 6X log2fold change at 1 year after the fall burn, but some species within *Massilia* had small (1-5) negative log2fold changes at 2 years after the fall and from 1 to 2 years after the spring burn.

**Fig. 8:**
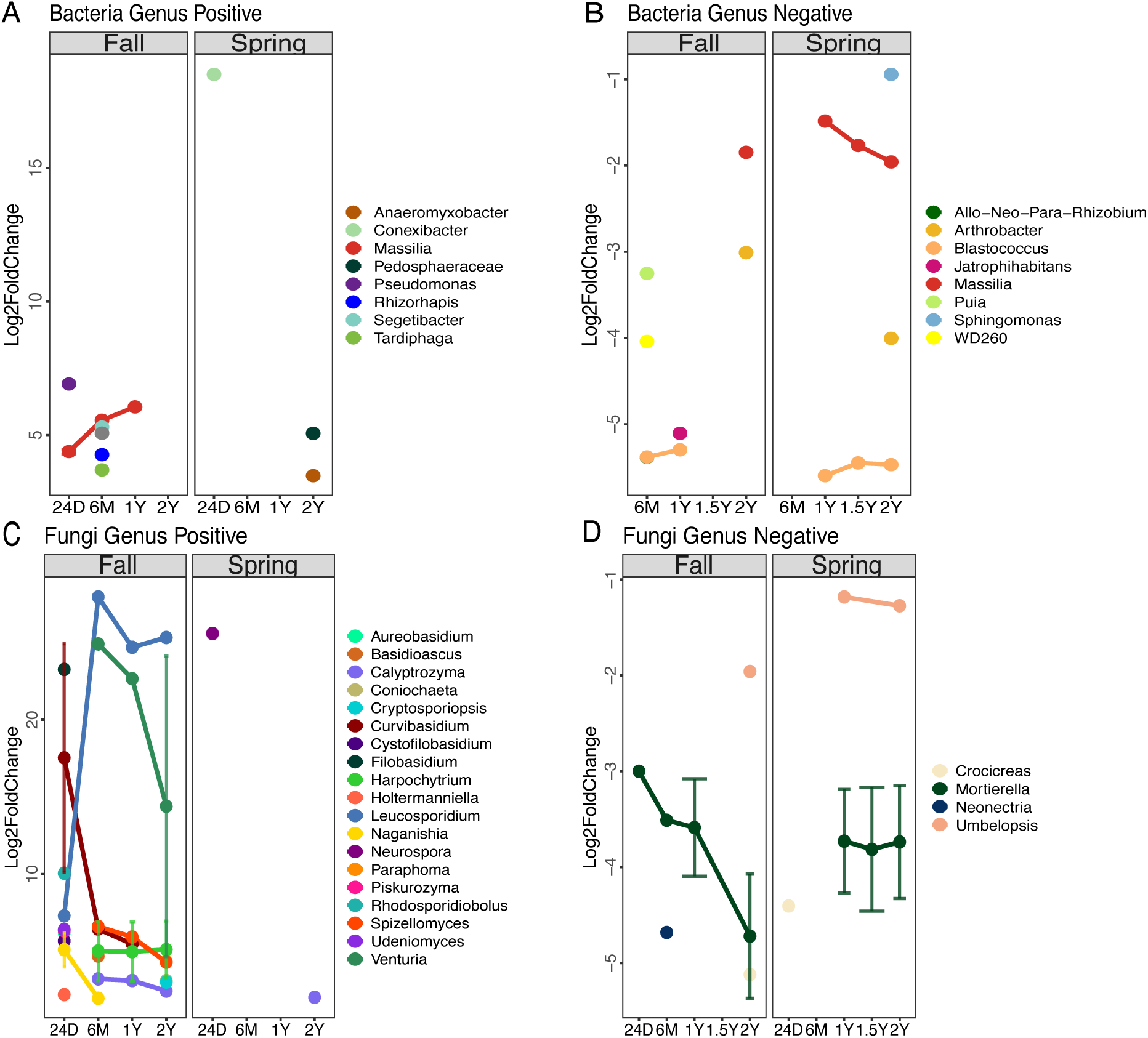
Summary figure of genera with DeSeq2 significant log2fold changes compared from each post-fire time point to pre-fire levels. Positive responders for A) bacteria and C) fungi, and negative responders for B) bacteria and D) fungi. Points representing taxa, and error bars represent the variability of species within the same genus.

For fungi, when testing significant changes in log2fold abundances in burned plots compared to pre-fire, 18 taxa in fall and 2 taxa in spring responded positively (Fig. 8C), whereas only 4 taxa in fall and 3 taxa in spring responded negatively (Fig. 8D). Overall, more fungal taxa had positive log2fold responses after the fall than spring burn. For example, Basidiomycota yeasts such as *Curvibasidium* (24 day to 1 year)*, Filobasidium* (24 days)*, Rhodosporidiobolus* (24 days), *Lecucosporidium* (24 days to 2 years), *Holtermanniella* (24 days), *Naganishia* (24 days to 6 months) and some Ascomycota such as *Venturia* (6 months to 2 years), *Harpocytrium* (6 months to 2 years), *Spizellomyces* (6 months to 2 years) positively changed with fall burn (Fig. 8C). After the spring burn, only 2 genera, both within the Ascomycota, *Neurospora* and *Calyptrozyma,* showed positive responses, with log2fold changes of 25X for *Neurospora* at 24 days and 2X for *Calyptrozyma* at 2 years (Fig. 8C).

## Discussion

Prescribed burns are increasingly used in both fall and spring seasons to restore fire-adapted ecosystems and mitigate the risk of severe wildfires. While prescribed burns typically have less deleterious effects on soil microbial communities than wildfires (Pressler et al., 2019), the impact of burn season on soil microbiomes was heretofore largely unknown. Here, we provide the first comprehensive test of fall versus spring burns on bacterial and fungal biomass, richness, and composition, and assess resilience across 2 years, in young montane coniferous forests. We found that, despite similar weather conditions during burns, Q1) fall burns reduced surface fuels more than spring burns and tended to produce more ash, which translated into fall burns having large negative impacts on Q2) bacterial and fungal biomass and richness, and Q3) EMF richness, whereas spring burns had no effect. Moreover, fall burns also resulted in Q4) more pronounced shifts in bacterial and fungal community composition, and more positively responding pyrophilous microbes. Although prescribed burns typically burn at much lower intensity than wildfires, our burns were hot enough to cause ∼29% (spring) to ∼36% (fall) tree mortality and led to the emergence of many taxa of pyrophilous microbes commonly found after severe wildfires for both fall and spring burns, including the bacterial genera *Massilia* and *Paenibacillus,* and fungal genera *Pyronema* and *Neurospora*. Despite having larger negative impacts in fall compared to spring burns, microbial biomass and richness nonetheless recovered within 2 years, although bacterial and fungal composition remained slightly altered at 2 years after both burns.

Although tree mortality and soil moisture remained consistent between fall and spring seasons in our study, higher fuel consumption (York et al., 2022) and deeper ash depth in fall burns indicated higher soil burn severity, which translated into larger reductions in microbial biomass and richness in fall than spring burns. We are unaware of any other study that comprehensively examined the impact of burn season on both bacteria and fungi. However, our results corroborate findings of greater impacts of fall compared to spring burns on AMF traits in grasslands (Hopkins & Bennett, 2023) and EMF richness in pine forests (Smith et al., 2004). Similarly, a study in a mixed conifer forest reported stronger effects of fall than spring burns on microbial respiration and enzyme activity (Hamman et al., 2008). Two other studies, which investigated the impacts of fall versus spring prescribed burns on fungi in Mediterranean Pine forests (Vázquez-Veloso et al., 2022) and Eastern Mediterranean shrublands (Livne-Luzon et al., 2021) found no impact of either burn on total fungal richness or abundance, likely because these burns were low intensity burns, whereas our prescribed fall burns were moderate intensity (York et al., 2022). Although they did not examine burn seasons, a smaller-scale study examining the impacts of a low-intensity prescribed burn (soil temperatures did not exceed 80°C) on soil bacteria and fungi in BFRS found that fungal but not bacterial richness declined in just one plot (Fischer et al., 2023).

While fall burns had more deleterious effects than spring burns on microbial biomass and richness, in all cases, microbial biomass and richness recovered within 2 years, suggesting strong microbial resilience after prescribed burns regardless of season. This contrasts sharply with the slow, decades-long recovery of bacteria and fungi seen after severe wildfires (Pérez-Valera et al., 2018). In our study, prescribed burns acted as a transient disturbance to soil microbial communities with limited long-term consequences. The rapid recovery of microbial biomass and richness highlights the benefits of prescribed burns to reduce fuel loads while preserving belowground ecological processes. Soil carbon dynamics mirrored these seasonal differences in burn severity (Krichels et al., 2026). Fall burns in these same plots combusted substantially more particulate and bulk soil carbon than spring burns without increasing pyrogenic organic matter, with carbon losses driven primarily by direct combustion and sustained microbial respiration that constrained post-fire carbon accumulation. Fall burns also had higher soil respiration immediately post-fire than spring burns in these same plots (Krichels et al., 2026), which is likely due to surviving microbes feasting on necromass from higher microbial mortality after fall burns (Buckeridge et al., 2022; Johnson et al., 2024). In contrast, spring burns were less severe, did not reduce particulate, mineral-associated, or pyrogenic carbon pools, and suppressed microbial respiration for over a year, coinciding with increases in mineral-associated and bulk soil carbon (Krichels et al., 2026). Together, these results indicate that while fall burns impose greater short-term microbial and carbon costs, spring burns better preserve soil carbon stocks while maintaining rapid microbial recovery.

We found that both fall and spring prescribed burns induced small but significant shifts in bacterial and fungal community composition, with stronger and more persistent effects following fall burns, indicating a shift in dominant taxa despite overall recovery in biomass and richness. In contrast, spring burns caused minimal changes in microbial composition, with biomass and richness largely unaffected. Similar compositional shifts without major losses in richness have also been observed following prescribed burns across diverse ecosystems, including grasslands (Glassman et al., 2023), boreal forests (Johnson et al., 2023), and the southern Appalachian Forest (Rafie et al., 2024), and also after a low-intensity desert fire (Joukhajian et al., 2026). These findings together suggest that while prescribed burns, particularly in the fall, may shift microbial community composition, they still support rapid recovery.

Despite relatively small changes in community composition compared to wildfires, our prescribed burns led to the emergence of several pyrophilous bacteria commonly seen after wildfires in both seasons. For example, Firmicutes increased in abundance after both fall and spring burns, similar to high-severity wildfires in redwood tanoak forests (Enright et al., 2022), chaparral shrublands (Pulido-Chavez et al., 2023), Colorado montane coniferous forests (Caiafa et al., 2023), and boreal coniferous forests (Whitman et al., 2019). In particular, the Firmicute genus *Paenibacillus,* which dominated our burned plots from 3 days to 1 year after the fall burn and at 3 days after the spring burn, also dominated after chaparral wildfire (Pulido-Chavez et al., 2023), likely due to its ability to produce thermotolerant endospores (Liu et al., 2014) and fix inorganic nitrogen (Liu et al., 2019). Post-fire changes in bacterial community composition were also driven by disappearances of certain groups like the Verrucromicrobiota, which was also negatively affected by wildfires in Colorado coniferous forests (Caiafa et al., 2023; Nelson et al., 2022). Additionally, the Proteobacteria genus *Massilia*, which is known to rapidly proliferate after wildfires due to its high 16S copy number (Johnson et al., 2023; Whitman et al., 2019) and putative ability to degrade PyOM (Pulido Barriga et al., 2025; Sari et al., 2026), dominated burned plots for up to 2 years in fall and up to 1.5 years in spring. While *Massilia* was consistently abundant across all plots prior to fire, DESeq2 analysis showed a positive response at 24 days after fall burns, followed by a slight decline two years later (Fig. 8A). Overall, our results indicate that while their increases in abundance are not as extreme as after wildfires in redwood tanoak forests (Enright et al., 2022) and chaparral shrublands (Pulido-Chavez et al., 2023), the fall burn still generated more pyrophilous bacteria than spring burns, consistent with higher soil burn severity in fall than spring.

While high-severity wildfires often cause fungi to shift in dominance from Basidiomycota to Ascomycota (Enright et al., 2022; Fox et al., 2022), here we found that Basidiomycota, mainly several genera of yeasts, increased in abundance after fall but not spring burns. For example, *Naganishia* increased in abundance after high-severity wildfires in Colorado montane coniferous forests (Caiafa et al., 2023) and in deserts (Canini et al., 2023; Joukhajian et al., 2026). *Geminibasidium* dominated here and after high-severity wildfires in chaparral shrublands (Pulido-Chavez et al., 2023) and redwood-tanoak forests (Enright et al., 2022). Further*, Geminibasidium* and *Solicoccozyma* increased in abundance after another prescribed burn in BFRS (Fischer et al., 2023) and *Solicoccozyma* after experimental pyrocosm burns from soils from a nearby Sierra Nevada pine forest (Bruns et al., 2020). Although there is no clear evidence correlating the basidiomycete yeast *Leucosporidium* to fire, it was highly abundant at 6 months after the fall burns, which was right after a snowstorm, which may have created an optimal environment for it to bloom since it has been reported in marine and Antarctic areas (Da Silva et al., 2024; Gonçalves et al., 2024). Among the few Ascomycota yeasts identified in our burned plots, *Calyptrozyma* exhibited the most significant changes in relative abundance after the fall burns, and this fungus also increased after high-severity wildfires in Washington Pine forest (Pulido-Chavez et al., 2021), Canadian boreal forests (Whitman et al., 2019), and a previous prescribed fire in BFRS (Fischer et al., 2023).

In addition to increases in Basidiomycota yeasts, changes in richness and composition after fall fires were likely driven by the loss of EMF, which, due to their symbiotic association with pines (Fujimura et al., 2005; Pulido-Chavez et al., 2021), were likely affected by the high tree mortality after fall burns. Like high severity wildfires in other EMF dominated systems (Caiafa et al., 2023; Pulido-Chavez et al., 2023), the EMF Basidiomycota genus *Inocybe* was abundant across time in our control plot but disappeared from our burned plots. Another Basidiomycota EMF genus, *Rhizopogon*, also showed higher abundance following fall burns, with particular increases in *Rhizopogon alkavirens*, consistent with the ecology of *Rhizopogon*, which is known to form persistent soil spore banks that can survive disturbance (Glassman et al., 2015). Previous studies from other pine forest soils showed that a different species within the genus, *Rhizopogon olivaceotinctus*, bloomed after high-severity wildfires (Baar et al., 1999; Glassman et al., 2016) and laboratory heat manipulations (Bruns et al., 2019; Peay et al., 2010). These suggest that fall burns significantly restructure EMF communities, with the loss of fire-sensitive taxa like *Inocybe*, while resilient spore-banking genera like *Rhizopogon* persist or increase, highlighting divergent post-fire recovery strategies among EMF.

Although the spring burns appeared to have minimal impacts on soil microbiomes, two well-known pyrophilous Ascomycota, *Pyronema* and *Neurospora*, temporarily bloomed in the first month after spring burns. While fire management typically aims for controlled and relatively uniform prescribed burns, like wildfires, the patchiness of the landscape will often result in heterogeneous burns that can lead to pyrodiversity (Hopkins & Bennett, 2023). Moreover, the uneven heat due to differential fuel load from the spring burn might trigger spore production (El-Abyad & Webster, 1968; Jalaluddin, 1967), enabling the temporary emergence of *Pyronema* and *Neurospora,* which were not similarly triggered by the hotter fall burn. Indeed, *Pyronema domesticum*, the first described pyrophilous fungus (Seaver, 1909), is known for rapid fruiting and mycelial proliferation after wildfires (Filialuna & Cripps, 2021; Pulido-Chavez et al., 2021; Pulido-Chavez et al., 2023) and experimental burns (Bruns et al., 2020). Recent genomic analysis showed that *Pyronema* likely dominates post-fire due to relatively rapid growth in the absence of competitors (Sari et al., 2026). Similarly, *Neurospora* has heat triggered spores (Emerson, 1948) and has also been reported to be abundant after low-intensity fires in desert environments (Joukhajian et al., 2026). This ephemeral but rapid response could explain why they were observed shortly after the spring burn but disappeared within a year, as the ecosystem quickly returned to a more stable state following the fire.

## Conclusion

While substantial evidence supports the use of prescribed burns to reduce fuel loads and maintain ecosystem health by mitigating aboveground wildfire impacts (Keeley & Brennan, 2012; Knapp et al., 2005; North et al., 2019; Ryan et al., 2013), implementing prescribed fire remains challenging due to logistic constraints, prompting the increased implementation of prescribed burns in spring outside of the historical burn window in summer and fall (Dong et al., 2022). In our study, fall prescribed burns significantly reduced bacterial and fungal biomass and richness, whereas spring burns did not. However, even after more severe fall burns, richness and biomass recovered within 2 years, which is much shorter than after high-severity wildfires (Pérez-Valera et al., 2018). Yet both fall and spring burns led to the emergence of pyrophilous taxa, which might contribute to overall ecosystem diversity. Since spring burns had subtle impacts on microbial communities, the use of spring burns may contribute to the overall ecosystem health by preserving microbial functions crucial for soil health and nutrient cycling, including the impacts on carbon sequestration (Krichels et al., 2026). Therefore, spring burns may be preferable when management objectives prioritize maintaining existing microbial community composition with minimal disturbance, whereas fall burns may be more suitable when maximizing fuel reduction is the primary goal, even if this entails greater but still resilient microbial compositional change. Overall, these findings provide a basis for developing adaptive, seasonally flexible prescribed fire regimes that meet ecological objectives while remaining operationally feasible, reinforcing prescribed fire as an effective and sustainable approach to ecological management.

## Data Availability Statement

The data supporting the findings of this study are available in the National Center for Biotechnology Information (NCBI) Sequence Read Archive under Bio-Project accession number PRJNA1417007; https://www.ncbi.nlm.nih.gov/sra/PRJNA1417007. All R scripts with statistical codes have been made available on GitHub https://github.com/Basubi1/Young-Blodgett-Rscript.

## Ethics Approval Statement

Not applicable (this study did not involve human participants or animals).

## Acknowledgments

This work was supported by the CALFIRE Forest Health Research Program Award 8GG21801 to SIG, RAY, and PMH. We thank Ariel Roughton and Amy Mason for aiding in permitting and site selection, and Hunter Noble for help with the prescribed burns. We thank BFRS for allowing us to sample. Thanks to Judy Chung for help with establishing the plots and pre-fire sampling, James Randolph for assistance with collecting and storing soil samples, and to Arik Joukhajian and Marcos Caiafa for advice in bioinformatics and statistical analysis. Thanks to anonymous reviewers for feedback on the manuscript.

## Author contributions

SIG and RAY conceived of the project idea and experimental design, SIG, RAY, and PMH obtained funding for the project, SIG and RAY set up the plots, RAY led the burns, RAY collected tree mortality and fire ecology data, BBZ performed soil processing, DNA extractions, and all molecular work, bioinformatics, and statistical analysis for the project, MFPC assisted in field work, bioinformatics, and statistical analysis, BBZ analyzed and interpreted the data and wrote the first draft, all authors contributed to editing of the manuscript.

## DATA AVAILABILITY STATEMENT

Raw sequence reads have been submitted to the National Centre for Biotechnology Information (NCBI) Sequence Read Archive under BioProject accession number PRJNA1417007. All R scripts with statistical codes have been made available on GitHub https://github.com/Basubi1/Young-Blodgett-Rscript.

