## supplementaltablesandfigures for "Differential impacts of fall versus spring prescribed burns on microbial biomass, richness, and composition in young mixed conifer forests"

MOLECULAR ECOLOGY

**Table of Contents:**


| | |
|---|---|
| Table S1: Experimental site description | Page 1 |
| Table S2: Moisture variation across plots (seasons), time and treatment | Page 2 |
| Table S3. Negative binomial generalized mixed effect models for bacteria | Page 4 |
| Table S4. Negative binomial generalized mixed effect models for bacteria | Page 6 |
| Table S5. Richness percent change for bacteria | Page 8 |
| Table S6. Richness percent change for fungi | Page 9 |
| Table S7. Fungal functional guild richness change | Page 10 |
| Table S8. PERMANOVA of bacterial and fungal community composition (time and treatment) | Page 12 |
| Table S9. Pairwise comparison (Adonis) of bacterial and fungal community composition | Page 13 |
| Figure S1: Fall vs spring tree mortality and Ash depth | Page 14 |
| Figure S2: Moisture variation across plots and time | Page 15 |
| Figure S3: Fall vs spring monthly average precipitation | Page 16 |
| Figure S4: Soil burn severity impact on bacterial and fungal richness | Page 17 |
| Figure S5: Soil burn severity impact on bacterial and fungal biomass | Page 18 |
| Figure S6: Relative abundance of fungal genera (< 2% percent sequence abundance) | Page 19 |

MOLECULAR ECOLOGY

**Table S1**. Blodgett Forest Research Station (BFRS) plot information. Blodgett Forest Research Station (BFRS) plot locations and elevations. We burned 4 plots in the fall (11/13/2019), 4 in the spring (4/28-5/1/2020), and 1 was left unburned (control), allowing us to have a complete Before-After-Control-Impact (BACI) design.

| Plot | Longitude | Latitude | Elevation (m) | Season |
|---|---|---|---|---|
| 50E | -120.66399 | 38.9136143 | 1322.73604 | Control |
| 260D | -120.66547 | 38.9110373 | 1337.1967 | Fall |
| 260E | -120.66546 | 38.909667 | 1342.68636 | Fall |
| 50G | -120.66929 | 38.918681 | 1282.34778 | Fall |
| 50H | -120.66961 | 38.914821 | 1304.83988 | Fall |
| 260F | -120.66584 | 38.9087017 | 1342.95032 | Spring |
| 260G | -120.66397 | 38.9080445 | 1330.16846 | Spring |
| 50I | -120.66748 | 38.9152677 | 1315.12553 | Spring |
| 50J | -120.66872 | 38.9134017 | 1323.31006 | Spring |

MOLECULAR ECOLOGY

**Table S2**. Soil moisture variation for all plots. Comparisons among a. all burned and control in fall versus spring plots, b. fall control versus fall burned, c. spring control versus spring burned, d. Fall pre-fire versus post-fire, e. Spring pre-fire versus post-fire. significance codes: '***' = P < 0.001, '**' = P < 0.01, based on the negative binomial generalized mixed effect models with plot, subplot, and collection day as random effects.

**a**.

| **All burned Fall vs Spring** | **Estimate** | **Std. Error** | **df** | **t value** | **Pr(>\|t\|)** |
|---|---|---|---|---|---|
| Intercept | 33.137 | 4.921 | 35.603 | 6.734 | 7.76e-08 *** |
| Fall:Spring: Control | 5.27 | 6. 05 | 112.00 | 0.87 | 0.38 |
| Fall:Spring: Prefire | 4.11 | 6. 05 | 112.00 | 0.77 | 0.24 |
| Fall:Spring:24D | 4.37 | 6. 05 | 112.00 | 0.72 | 0.47 |
| Fall:Spring:1Y | -6.58 | 6. 05 | 112.00 | -1.08 | 0.27 |
| Fall:Spring:2Y | -5.34 | 6.05 | 112 | -1.00 | 0.40 |

**b**.

| **Fall control vs Fall burned** | **Estimate** | **Std. Error** | **df** | **t value** | **Pr(>\|t\|)** |
|---|---|---|---|---|---|
| Fall Control: 24D | 9.65 | 2.29 | 137.00 | 4.20 | 4.66 |
| Fall Control: 6M | 12.93 | 5.13 | 137.00 | 2.52 | 0.01 ** |
| Fall Control: 1Y | 9.92 | 5.13 | 137.00 | 1.93 | 0.05 * |
| Fall Control: 2Y | 9.38 | 5.13 | 137.00 | 1.83 | 0.06 |

**c**.

| **Spring control vs Spring burned** | **Estimate** | **Std. Error** | **df** | **t value** | **Pr(>\|t\|)** |
|---|---|---|---|---|---|
| Spring Control: 3D | 8.70 | 5.48 | 165.00 | 1.58 | 0.11 |
| Spring Control: 24D | 18.99 | 5.48 | 165.00 | 3.46 | 0.001*** |
| Spring Control: 1Y | 6.47 | 5.48 | 165.00 | 1.18 | 0.23 |
| Spring Control: 1.5Y | 8.01 | 5.48 | 165.00 | 1.46 | 0.14 |
| Spring Control: 2Y | 14.15 | 5.48 | 165.00 | 2.57 | 0.01 ** |

**d**.

| **Fall pre-vs post-fire)** | **Estimate** | **Std. Error** | **df** | **t value.** | **Pr(>\|t\|)** |
|---|---|---|---|---|---|
| (Intercept) | 33.52 | 2.88 | 6.44 | 11.61 | 1.47e-05*** |

MOLECULAR ECOLOGY

| | | | | | |
|---|---|---|---|---|---|
| Prefire: 24D | 9.52 | 2.10 | 141.00 | 4.51 | 1.32e-05*** |
| Prefire: 6M | 0.75 | 2.10 | 141.00 | 0.35 | 0.72 |
| Prefire: 1Y | 0.67 | 2.10 | 141.00 | 0.32 | 0.74 |
| Prefire: 2Y | 7.15 | 2.10 | 141.00 | 3.39 | 0.001 *** |

**e**.

| **Spring pre- vs post-fire)** | **Estimate** | **Std. Error** | **df** | **t value.** | **Pr(>\|t\|)** |
|---|---|---|---|---|---|
| (Intercept) | 33.84 | 2.72 | 7.81 | 12.44 | 1.98e-06 *** |
| Prefire: 3D | 4.13 | 2.25 | 170.00 | 1.83 | 0.06 |
| Prefire: 24D | -0.47 | 2.25 | 170.00 | -0.21 | 0.83 |
| Prefire: 1Y | 2.05 | 2.25 | 170.00 | 0.90 | 0.36 |
| Prefire: 1.5Y | 8.25 | 2.25 | 170.00 | 3.66 | 0.0003 *** |

MOLECULAR ECOLOGY

**Table S3.** Effects of burn season on bacterial richness. Generalized linear mixed models (GLMMs) results based on the negative binomial distribution with plot, subplot, and collection day as random effects and treatment (pre- versus post-fire) and monthly average precipitation (mm) as fixed effects. Significance codes: '***' = P < 0.001, '**' = P < 0.01.

a. Treatment: Fall Burn (Bacteria)

| Fixed effects: | Estimate | Std. Error | z value | Pr(>\|z\|) |
|---|---|---|---|---|
| (Intercept) | 5.93 | 0.11 | 51.00 | < 2e-16 *** |
| 3D-PostFire. | -0.15 | 0.12 | -1.19 | 0.23 |
| 24D-PostFire. | -0.41 | 0.12 | -3.29 | 0.001 ** |
| 6M-PostFire. | -0.38 | 0.12 | -3.11 | 0.009 *** |
| 1Y-PostFire | -0.33 | 0.12 | -2.67 | 0.007 ** |
| 2Y-PostFire | -0.20 | 0.12 | -0.46 | 0.06 |
| Precipitation | 0.021 | 0.45 | 0.05 | 0.96 |

b. Treatment: Fall Control (Bacteria)

| Fixed effects: | Estimate | Std. Error | z value | Pr(>\|z\|) |
|---|---|---|---|---|
| (Intercept) | 5.86 | 0.06 | 87.78 | < 2e-16 *** |
| 3D-PostFire | 0.48 | 0.09 | 5.16 | 2.47e-07 *** |
| 24D-PostFire | 0.13 | 0.09 | 1.46 | 0.14 |
| 6M-PostFire | 0.26 | 0.09 | 1.87 | 0.06 |
| 1Y-PostFire | 0.23 | 0.09 | 1.48 | 0.14 |
| 2Y-PostFire | 0.03 | 0.09 | 0.41 | 0.67 |
| Precipitation | 0.022 | 0.40 | 0.04 | 0.93 |

c. Treatment – Spring burn (Bacteria)

| Fixed effects: | Estimate | Std. Error | z value | Pr(>\|z\|) |
|---|---|---|---|---|
| (Intercept) | 6.05 | 0.05 | 115.23 | < 2e-16 *** |
| 3D-PostFire | 0.02 | 0.06 | 0.38 | 0.69 |
| 24D-PostFire | 0.04 | 0.06 | 0.73 | 0.46 |
| 1Y-PostFire | -0.01 | 0.06 | -1.66 | 0.09 |
| 1.5Y-PostFire | 0.03 | 0.06 | 0.51 | 0.61 |

MOLECULAR ECOLOGY

| | | | | |
|---|---|---|---|---|
| 2Y-PostFire | 0.06 | 0.06 | 1.05 | 0.23 |

d. Treatment- Spring Control (Bacteria)

| Fixed effects: | Estimate | Std. Error | z value | Pr(>\|z\|) |
|---|---|---|---|---|
| (Intercept) | 5.86 | 0.06 | 95.49 | <2e-16 *** |
| 3D-PostFire | 0.26 | 0.08 | 3.12 | 0.06 |
| 24D-PostFire | 0.14 | 0.08 | 1.71 | 0.08 |
| 1Y-PostFire | 0.10 | 0.08 | 1.18 | 0.23 |
| 1.5Y-PostFire | 0.03 | 0.08 | 0.45 | 0.65 |
| 2Y-PostFire | 0.21 | 0.08 | 2.52 | 0.06 |

MOLECULAR ECOLOGY

**Table S4.** Effects of burn season on fungal richness. Generalized linear mixed models (GLMMs) results based on the negative binomial distribution with plot, subplot, and collection day as random effects and treatment (pre- versus post-fire) as fixed effect, and monthly average precipitation (mm). Significance codes: '***' = P < 0.001, '**' = P < 0.01.

a. Treatment – Fall burn (Fungi)

| Fixed effects | Estimate | Std. Error | z value | Pr(>\|z\|) |
|---|---|---|---|---|
| (Intercept) | 5.38 | 0.09 | 56.09 | < 2e-16 *** |
| 3D-PostFire | -0.29 | 0.10 | -2.84 | 0.004 ** |
| 24D-PostFire | -0.29 | 0.10 | -2.89 | 0.004 ** |
| 6M-PostFire | -0.38 | 0.10 | -3.77 | 0.0001*** |
| 1Y-PostFire | -0.17 | 0.10 | -1.69 | 0.09 |
| 2Y-PostFire | -0.66 | 0.10 | -1.79 | 0.07 |
| Precipitation | 0.52 | 0.37 | 1.40 | 0.16 |

b. Treatment – Fall Control (Fungi)

| Fixed effects: | Estimate | Std. Error | z value | Pr(>\|z\|) |
|---|---|---|---|---|
| (Intercept) | 5.60 | 0.06 | 90.02 | <2e-1.6 *** |
| 3D-PostFire | -0.03 | 0.08 | -0.37 | 0.70 |
| 24D-PostFire | 0.01 | 0.08 | 0.15 | 0.87 |
| 6M-PostFire | 0.07 | 0.08 | 0.79 | 0.42 |
| 1Y-PostFire | 0.08 | 0.08 | 1.00 | 0.31 |
| 2Y-PostFire | 0.08 | 0.08 | 1.00 | 0.03 |
| Precipitation | 0.50 | 0.27 | 1.40 | 0.09 |

c. Treatment – Spring burn (Fungi)

| Fixed effects: | Estimate | Std. Error | z value | Pr(>\|z\|) |
|---|---|---|---|---|
| (Intercept) | 5.57 | 0.05 | 96.62 | < 2e-16 *** |
| 3D-PostFire | -0.02 | 0.06 | -0.28 | 0.77 |
| 24D-PostFire | -0.12 | 0.06 | -1.77 | 0.07 |

| | | | | |
|---|---|---|---|---|
| 1Y-PostFire | 0.01 | 0.06 | 0.14 | 0.88 |
| 1.5Y-PostFire | 0.03 | 0.06 | 0.53 | 0.59 |
| 2Y-PostFire | 0.02 | 0.06 | 0.55 | 0.66 |
| Precipitation | -0.00 | 0.00 | -0.75 | 0.45 |

d. Treatment- Control (Fungi)

| Fixed effects: | Estimate | Std. Error | z value | Pr(>\|z\|) |
|---|---|---|---|---|
| (Intercept) | 5.60 | 0.05 | 95.12 | <2e-16 *** |
| 3D-PostFire | 0.07 | 0.08 | 0.84 | 0.39 |
| 24D-PostFire | 0.02 | 0.08 | 0.27 | 0.78 |
| 1Y-PostFire | 0.01 | 0.08 | 0.14 | 0.88 |
| 1.5Y-PostFire | 0.18 | 0.08 | 2.27 | 0.06 |
| 2Y-PostFire | 0.02 | 0.08 | 0.35 | 0.72 |
| Precipitation | 0.60 | 0.35 | 1.40 | 0.06 |

**Table S5:** Bacterial richness percent change from pre-fire in burned plots in both seasons, with means based on 6 subsamples per plot at each time point.

| Season | Time | Richness means per subplot | Standard Deviation (SD) | Standard error of the mean (SE) | Richness percent change |
|---|---|---|---|---|---|
| Fall | Pre-Fire | 503.87 | 207.92 | 42.44 | |
| Fall | 3D | 447.77 | 155.91 | 33.24 | -11.13 |
| Fall | 24D | 352 | 256.82 | 52.42 | -30.14 |
| Fall | 6M | 343.04 | 167.64 | 34.22 | -31.91 |
| Fall | 1Y | 354.26 | 87.1 | 18.17 | -29.69 |
| Fall | 2Y | 412.87 | 115.68 | 23.61 | -18.06 |
| Spring | Pre-Fire | 392.70 | 90.51 | 18.47 | |
| Spring | 3D | 402.62 | 96.15 | 19.62 | 2.46 |
| Spring | 24D | 408 | 63.27 | 12.91 | 3.74 |
| Spring | 1Y | 359.65 | 116.69 | 24.33 | -8.41 |
| Spring | 1.5Y | 403.83 | 72.44 | 14.78 | 2.83 |
| Spring | 2Y | 417.5 | 87.74 | 17.91 | 6.31 |

MOLECULAR ECOLOGY

**Table S6:** Fungal percent change from pre-fire in burned plots in both seasons, with means based on 6 subsamples per plot at each time point.

| Season | Time | Richness means per subplot | SD | SE | Richness percent change |
|---|---|---|---|---|---|
| Fall | PreFire | 273.29 | 37.07 | 7.56 | |
| Fall | 3D | 213.59 | 71.46 | 15.23 | -21.84 |
| Fall | 24D | 208.83 | 67.040 | 13.68 | -23.58 |
| Fall | 6M | 191.5 | 84.47 | 17.24 | -29.92 |
| Fall | 1Y | 235.30 | 73.25 | 15.27 | -13.89 |
| Fall | 2Y | 240.83 | 93.52 | 19.09 | -11.87 |
| Spring | PreFire | 277.5 | 40.62 | 8.29 | |
| Spring | 3D | 273.36 | 90.42 | 20.74 | -1.48 |
| Spring | 24D | 245.80 | 57.70 | 12.59 | -11.42 |
| Spring | 1Y | 283.54 | 99.01 | 20.21 | 2.17 |
| Spring | 1.5Y | 282.72 | 65.74 | 14.01 | 1.88 |
| Spring | 2Y | 264.5 | 48.29 | 10.29 | -4.68 |

**Table S7:** Impact of fire on fungal guild richness: ectomycorrhizal fungi (EMF), saprobe, and plant pathogens. Percent change in burned plots compared to pre-fire in both seasons, based on 6 subsamples per plot at each time point.

| **Guilds** | **Season** | **Time** | **Mean Per Subplot** | **SD** | SE | **Percent change** |
|---|---|---|---|---|---|---|
| EMF | Fall | PreFire | 13.08 | 8.48 | 1.74 | |
| EMF | Fall | 3D | 13.23 | 8.64 | 1.84 | 1.10 |
| EMF | Fall | 24D | 7.13 | 6.18 | 1.26 | -45.54 |
| EMF | Fall | 6M | 6.88 | 5.31 | 1.08 | -47.45 |
| EMF | Fall | 1Y | 8.63 | 6.03 | 1.23 | -34.08 |
| EMF | Fall | 2Y | 11.58 | 5.16 | 1.05 | -11.46 |
| EMF | Spring | PreFire | 10.67 | 4.79 | 0.77 | |
| EMF | Spring | 3D | 9.81 | 6.91 | 0.75 | -8.04 |
| EMF | Spring | 24D | 8.14 | 3.44 | 0.75 | -23.66 |
| EMF | Spring | 1Y | 9.79 | 5.85 | 0.98 | -8.20 |
| EMF | Spring | 1.5Y | 9.75 | 3.77 | 1.19 | -8.59 |
| EMF | Spring | 2Y | 9.04 | 3.67 | 1.51 | -15.23 |
| Saprobe | Fall | PreFire | 95.75 | 16.61 | 3.39 | 0.00 |
| Saprobe | Fall | 3D | 76.82 | 25.32 | 5.40 | -19.77 |
| Saprobe | Fall | 24D | 69.38 | 28.54 | 5.83 | -27.55 |
| Saprobe | Fall | 6M | 64.54 | 31.31 | 6.39 | -32.59 |
| Saprobe | Fall | 1Y | 71.96 | 30.58 | 6.24 | -24.85 |
| Saprobe | Fall | 2Y | 76.33 | 35.28 | 7.20 | -20.28 |
| Saprobe | Spring | PreFire | 95.25 | 19.55 | 3.99 | 0.00 |
| Saprobe | Spring | 3D | 87.14 | 43.80 | 9.56 | -8.51 |
| Saprobe | Spring | 24D | 78.57 | 24.01 | 5.24 | -17.51 |
| Saprobe | Spring | 1Y | 92.25 | 43.10 | 8.80 | -3.15 |
| Saprobe | Spring | 1.5Y | 87.54 | 27.51 | 5.62 | -8.09 |
| Saprobe | Spring | 2Y | 83.33 | 26.75 | 5.46 | -12.51 |
| PlantPathogen | Fall | PreFire | 8.54 | 2.55 | 0.52 | 0.00 |
| PlantPathogen | Fall | 3D | 6.27 | 3.28 | 0.70 | -26.56 |
| PlantPathogen | Fall | 24D | 6.50 | 3.16 | 0.65 | -23.90 |

| | | | | | | |
|---|---|---|---|---|---|---|
| PlantPathogen | Fall | 6M | 5.63 | 3.06 | 0.63 | -34.15 |
| PlantPathogen | Fall | 1Y | 6.79 | 3.26 | 0.66 | -20.49 |
| PlantPathogen | Fall | 2Y | 8.21 | 4.02 | 0.82 | -3.90 |
| PlantPathogen | Spring | PreFire | 9.46 | 2.70 | 0.55 | 0.00 |
| PlantPathogen | Spring | 3D | 8.05 | 5.32 | 1.16 | -14.92 |
| PlantPathogen | Spring | 24D | 6.38 | 4.18 | 0.91 | -32.54 |
| PlantPathogen | Spring | 1Y | 9.67 | 4.35 | 0.89 | 2.20 |
| PlantPathogen | Spring | 1.5Y | 10.33 | 4.04 | 0.82 | 9.25 |
| PlantPathogen | Spring | 2Y | 9.50 | 4.75 | 0.97 | 0.44 |

MOLECULAR ECOLOGY

**Table S8.** Impact of burn season and time on microbial community composition. We present statistics from permutational multivariate analysis of variance (PERMANOVA) of bacterial and fungal community composition response variable against effects of treatment (pre vs post-fire), time, and their interactions.

| Variables | | Fall | | | | Spring | | | |
|---|---|---|---|---|---|---|---|---|---|
| | | **Sum of Sqs.** | **$R^2$** | **F** | **P value** | **Sum of Sqs.** | **$R^2$** | **F** | **P value** |
| Bacteria | Fire | 1.33 | 0.03 | 4.57 | 0.0001 | 0.60 | 0.02 | 2.47 | 0.0001 |
| | Time | 3.47 | 0.09 | 2.45 | 0.0001 | 2.92 | 0.08 | 2.50 | 0.0001 |
| | Fire*Time | 3.49 | 0.08 | 2.77 | 0.0001 | 5.03 | 0.09 | 2.32 | 0.0001 |
| Fungi | Fire | 1.36 | 0.03 | 4.34 | 0.0001 | 0.88 | 0.02 | 2.90 | 0.0001 |
| | Time | 5.29 | 0.11 | 4.21 | 0.0001 | 4.33 | 0.11 | 3.03 | 0.0001 |
| | Fire*Time | 6.42 | 0.15 | 4.77 | 0.0001 | 4.62 | 0.15 | 4.28 | 0.0001 |

MOLECULAR ECOLOGY

**Table S9**. Pairwise comparison (adonis) for microbial community composition turnover at each time point, comparing either the burned plots or the unburned control plots to pre-fire. We used an adjusted p-value based on multiple pairwise comparisons (PERMANOVA). Significant turnovers are bolded.

| Pairs | P-value Fall Burned | P-value Fall Control | P-value Spring Burned | P-value Spring Control |
|---|---|---|---|---|
| Bacteria | | | | |
| PreFire vs 3D-PostFire | **0.01** | 0.12 | **0.01** | 0.67 |
| PreFire vs 24D-PostFire | **0.005** | 0.06 | **0.01** | **0.04** |
| PreFire vs 6M-PostFire (fall)&<br>PreFire vs 1Y-PostFire (Spring) | **0.005** | **0.01** | **0.01** | 0.09 |
| PreFire vs 1Y-PostFire (Fall)&<br>PreFire vs 1.5Y-PostFire (Spring) | **0.005** | **0.05** | **0.04** | **0.03** |
| PreFire vs 2Y-PostFire | **0.005** | **0.01** | **0.01** | 0.06 |

| Fungi | | | | |
|---|---|---|---|---|
| PreFire vs 3D-PostFire | 0.24 | **0.04** | **0.01** | 0.06 |
| PreFire vs 24D-PostFire | **0.01** | **0.03** | **0.01** | 0.06 |
| PreFire vs 6M-PostFire (fall)<br>PreFire vs 1Y-PostFire (Spring) | **0.01** | 0.09 | **0.03** | 0.06 |
| PreFire vs 1Y-PostFire (Fall)<br>PreFire vs 1.5Y-PostFire (Spring) | **0.03** | 0.07 | **0.01** | 0.06 |
| PreFire vs 2Y-PostFire | **0.01** | **0.04** | **0. 01** | **0.04** |

MOLECULAR ECOLOGY

**Figure S1:** Box plots showing mean plus or minus standard error of mean for A. Cumulative percent tree mortality at one year after fire and B. Ash depth in cm at 3 days after fall (yellow) versus spring burns (blue).

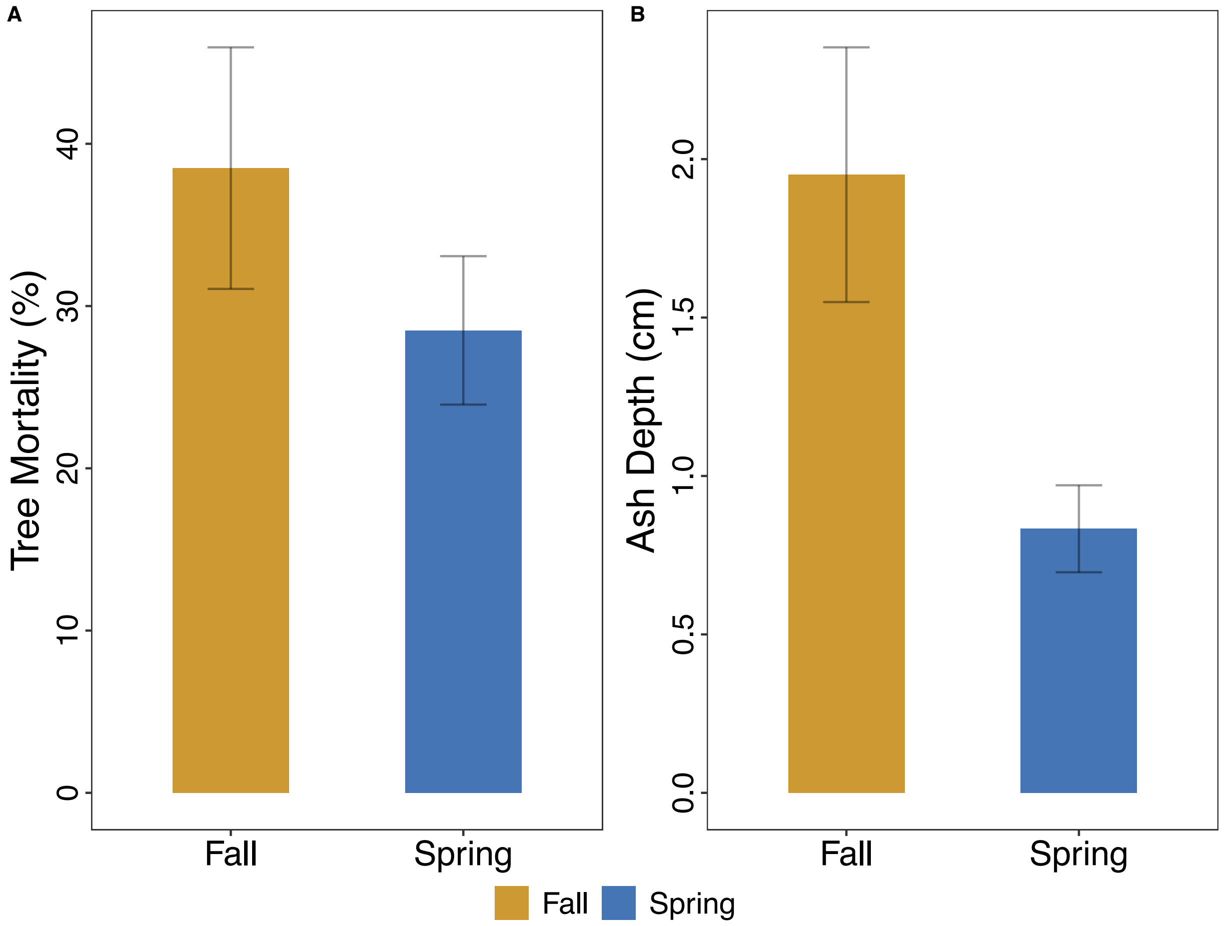

MOLECULAR ECOLOGY

**Figure S2:** Gravimetric percent soil moisture showing mean of 6 subsamples per plus or minus standard error of mean in fall versus spring burns in burned vs control plots. Shapes indicating Fall (triangle), Spring (square), versus Control (circle) plots, and blue indicates control or pre-fire, whereas red indicates burned plots post-fire. Asterisks indicate significant differences across time comparing each post-fire timepoint to pre-fire for either control (blue asterisks) and burned (red asterisks) samples with the following significance codes: '***' = P < 0.001, '**' = P < 0.01. Asterisks do not provide a direct comparison of burned vs control plots.

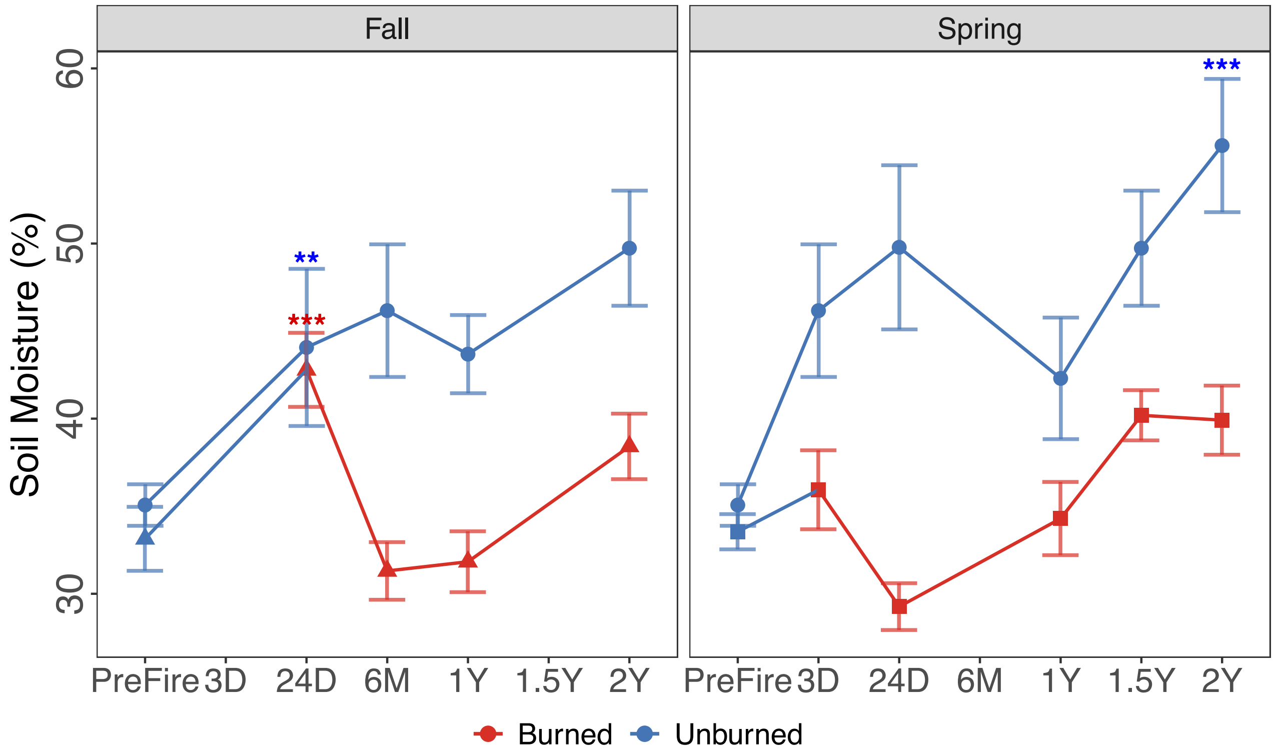

MOLECULAR ECOLOGY

**Figure S3:** Monthly average Precipitation (mm), corresponding to each timepoint when the samples were collected.

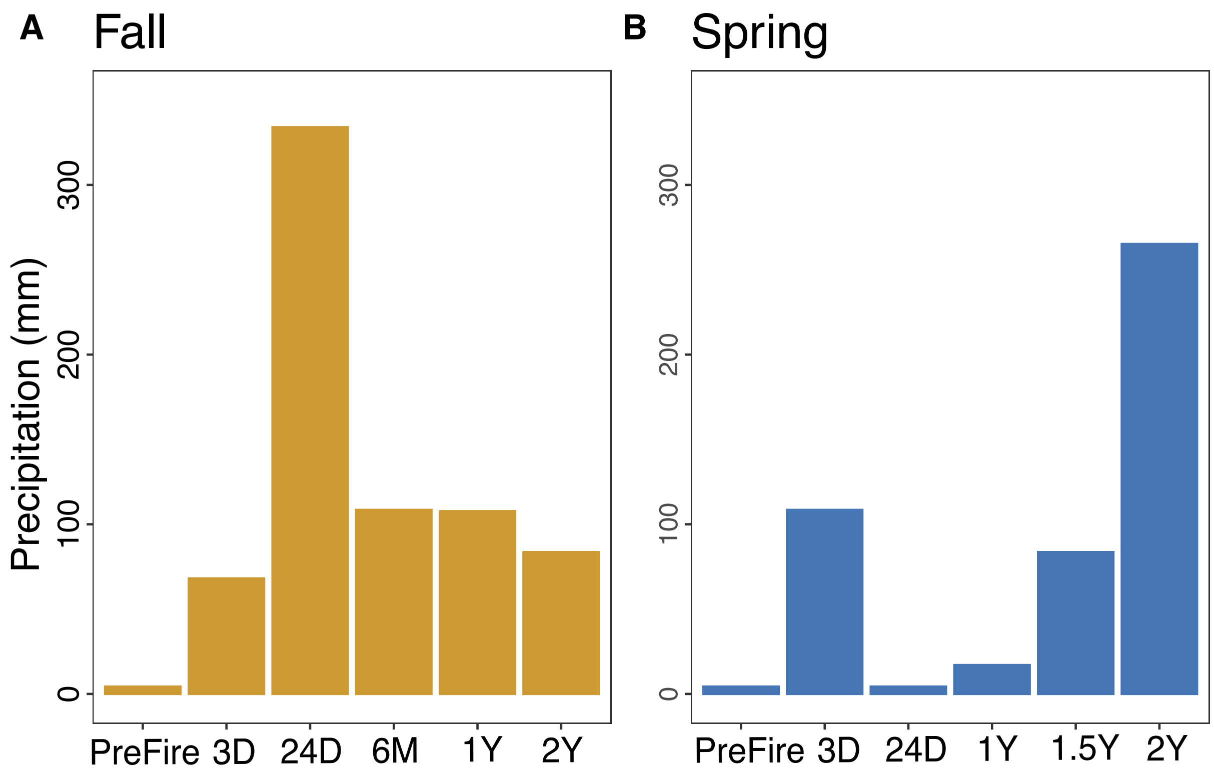

MOLECULAR ECOLOGY

**Fig S4:** Impact of soil burn severity on bacterial and fungal richness in fall versus spring. Bacterial richness in A) fall versus B) spring and fungal richness in C) fall versus D) spring regressed against ash depth in cm at 3 days post-fire as a proxy of soil burn severity. The blueline represents the model's prediction based on negative binomial regression, the grey is the standard error, and the p-value and adjusted $R^2$ value are listed in the top right corner.

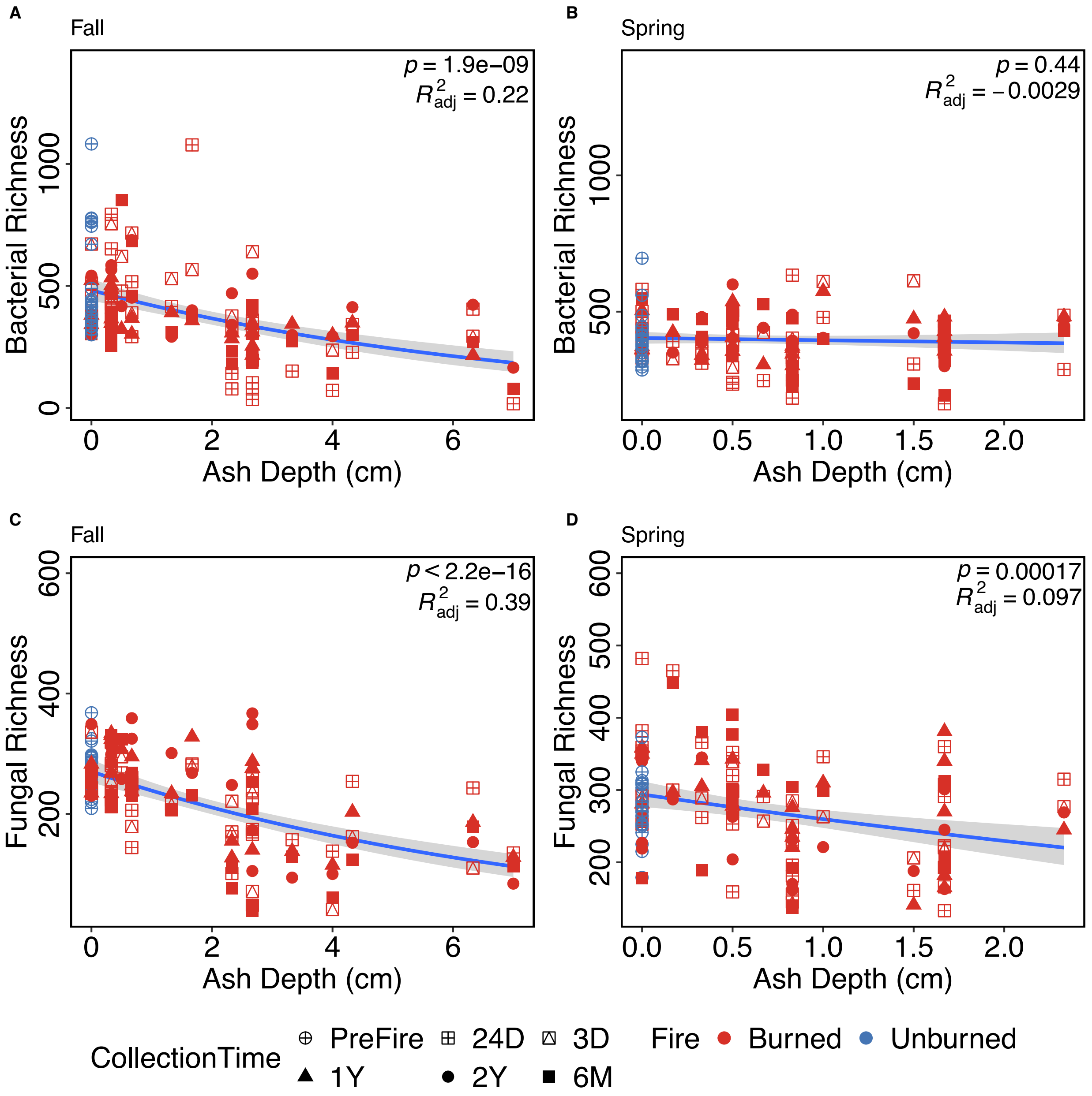

MOLECULAR ECOLOGY

**Fig S5:** Impact of soil burn severity on bacterial and fungal abundance as estimated by gene copy number in fall versus spring. Bacterial 16S copy number in A) fall versus B) spring and fungal 18S copy number in C) fall versus D) spring regressed against ash depth in cm at 3 days post-fire as a proxy of soil burn severity. The blueline represents the model's prediction based on negative binomial regression, the grey is the standard error, and the p-value and adjusted $R^2$ value are listed in the top right corner.

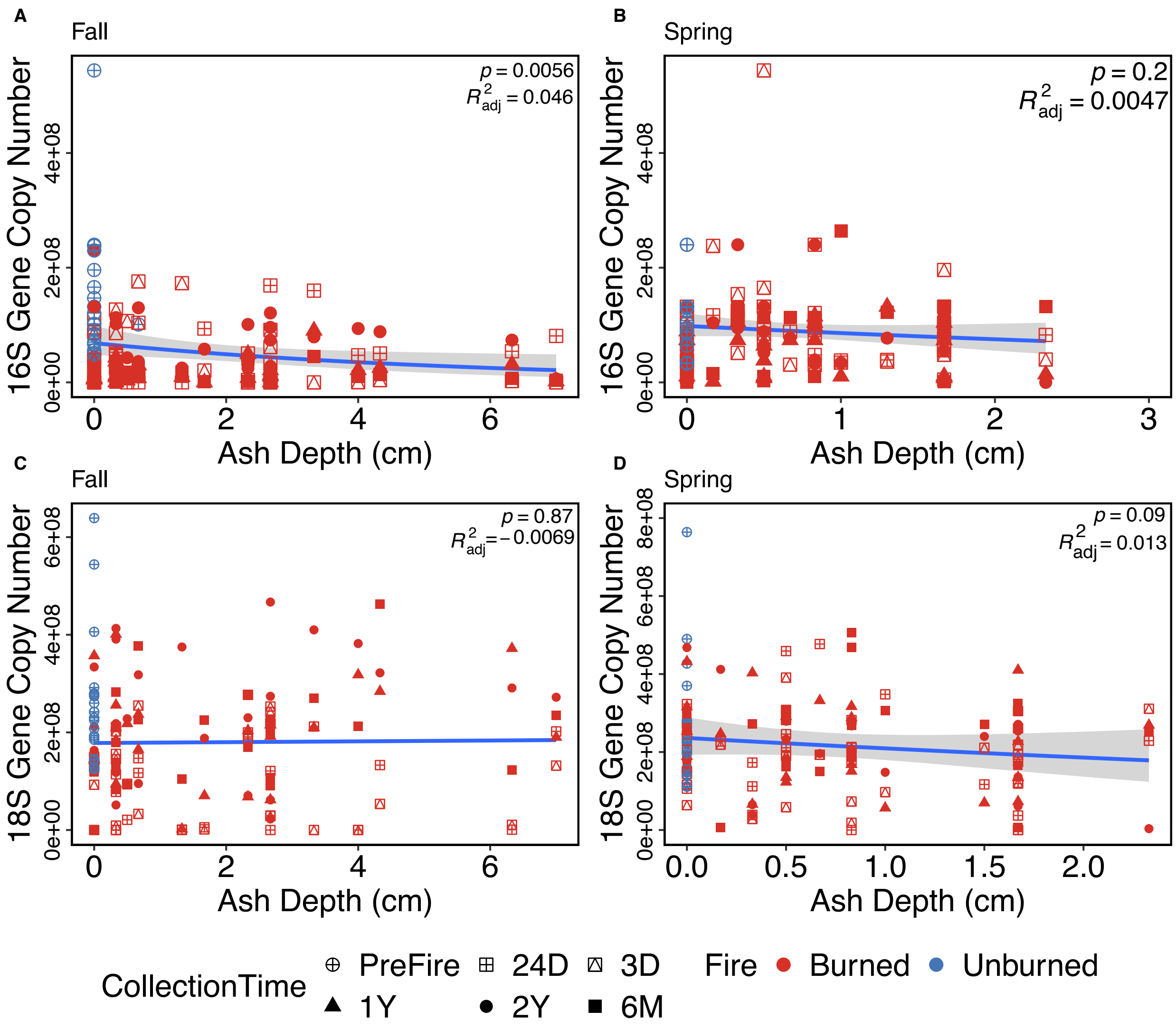

MOLECULAR ECOLOGY

**Fig S6:** Relative percent sequence abundance of fungal genera in the burned plots for A) fall and B) spring at each time point. Pre-fire and 3 days (3D), 24 days (24 D), 6 months (6M), 1 year (1 year), 1.5 years for the spring plots (1.5 Y), and 2 years (2Y). All phyla that were less than 2% abundant are condensed to improve visualization.

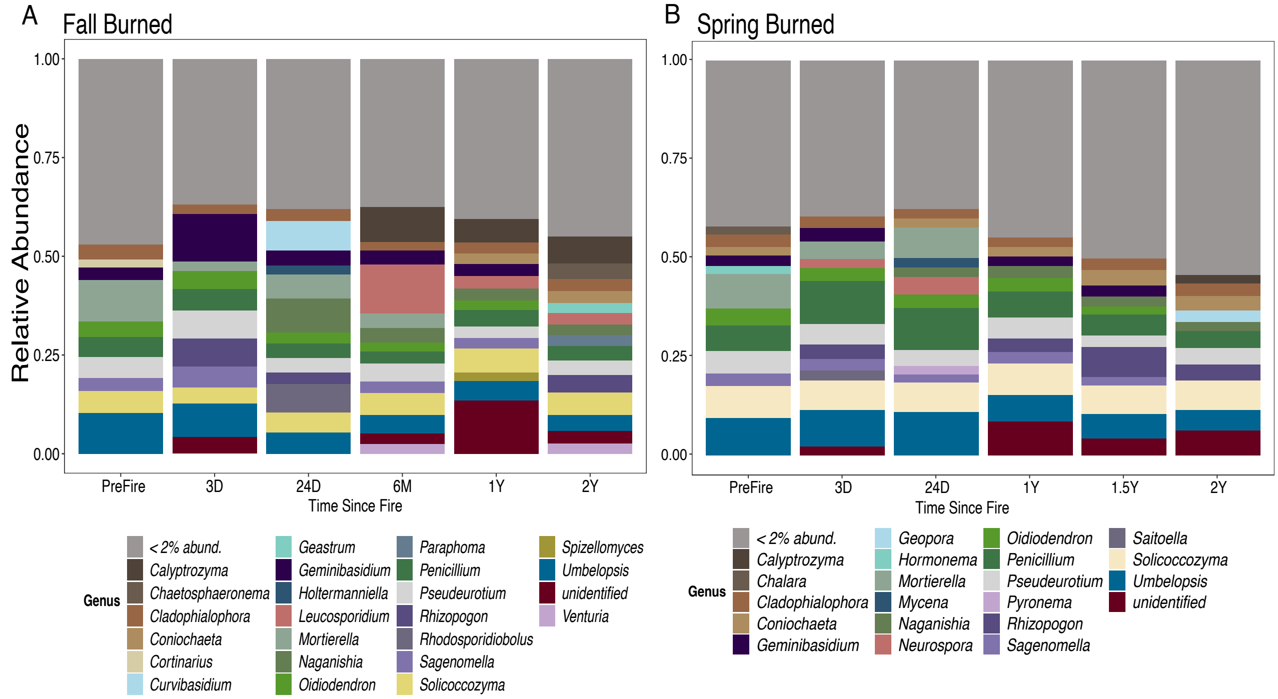
